# RiboTag translatomic profiling of *Drosophila* oenocytes under aging and oxidative stress

**DOI:** 10.1101/272179

**Authors:** Kerui Huang, Wenhao Chen, Fang Zhu, Hua Bai

## Abstract

**Background:** Aging is accompanied with loss of tissue homeostasis and accumulation of cellular damages. As one of the important metabolic centers, aged liver shows altered lipid metabolism, impaired detoxification pathway, increased inflammation and oxidative stress response. However, the mechanisms for these age-related changes still remain unclear. In fruit flies, *Drosophila melanogaster*, liver-like functions are controlled by two distinct tissues, fat body and oenocytes. Although the role of fat body in aging regulation has been well studied, little is known about how oenocytes age and what are their roles in aging regulation. To address these questions, we used cell-type-specific ribosome profiling (RiboTag) to study the impacts of aging and oxidative stress on oenocyte translatome in *Drosophila*.

**Results:** We show that aging and oxidant paraquat significantly increased the levels of reactive oxygen species (ROS) in adult oenocytes of *Drosophila*, and aged oenocytes exhibited reduced sensitivity to paraquat treatment. Through RiboTag sequencing, we identified 3324 and 949 differentially expressed genes in oenocytes under aging and paraquat treatment, respectively. Aging and paraquat exhibit both shared and distinct regulations on oenocyte translatome. Among all age-regulated genes, mitochondrial, proteasome, peroxisome, fatty acid metabolism, and cytochrome P450 pathways were down-regulated, whereas DNA replication and glutathione metabolic pathways were up-regulated. Interestingly, most of the peroxisomal genes were down-regulated in aged oenocytes, including peroxisomal biogenesis factors and beta-oxidation genes. Further analysis of the oenocyte translatome showed that oenocytes highly expressed genes involving in liver-like processes (e.g., ketogenesis). Many age-related transcriptional changes in oenocytes are similar to aging liver, including up-regulation of Ras/MAPK signaling pathway and down-regulation of peroxisome and fatty acid metabolism.

**Conclusions:** Our oenocyte-specific translatome analysis identified many genes and pathways that are shared between *Drosophila* oenocytes and mammalian liver, highlighting the molecular and functional similarities between the two tissues. Many of these genes are altered in both aged oenocytes and aged liver, suggesting a conserved molecular mechanism underlying oenocyte and liver aging. Thus, our translatome analysis will contribute significantly to the understanding of oenocyte biology, and its role in lipid metabolism, stress response and aging regulation.

## Introduction

Aging is the major risk factor for many chronic diseases [1]. The prevalence of liver diseases, such as non-alcoholic fatty liver disease (NAFLD), increase dramatically in the elderly [2, 3]. It is known that aging is associated with alterations of hepatic structure, physiology and function [4]. For example, aged liver shows reduced blood flow, loss of regenerative capacity, decreases in detoxification and microsomal proteins synthesis, increases in polyploidy, oxidative stress and mitochondrial damage [5]. Additionally, the metabolism for low-density lipoprotein cholesterol decreases by 35% [3]. Age-related increases in neutral fat levels and high-density lipoprotein cholesterol predispose aged liver to NAFLD and other liver diseases. Accumulated evidence suggests that age-related decline of liver function can be attributed to increased ROS production, DNA damage, activation of p300-C/EBP-dependent neutral fat synthesis [6], decreases in autophagy, increases in inflammatory responses [7, 8], and activation of nuclear factor-κB (NF-κB) pathway [4, 9]. Despite the genetic and functional analysis of liver aging and liver diseases, only a few studies have looked at the global transcriptional changes during liver aging [10-12].

Similar to mammals, the fruit fly (*Drosophila melanogaster*, hereafter as *Drosophila*) also shows age-dependent decline of tissue function and loss of homeostasis (reviewed in [13]). In *Drosophila*, liver-like functions are shared by two distinct tissues, fat body and oenocytes [14]. Fat body is the main tissue for energy storage in insects, and it plays a key role in metabolism, nutrition sensing, growth and immunity (reviewed in [15]). Fat body has also been implicated in the regulation of organismal aging [16]. Many longevity pathways act on fat body to control lifespan [17-19]. Compared to fat body, little is known about how oenocytes age and what is the role of oenocytes in aging regulation. Oenocytes are specialized hepatocyte-like cells responsible for energy metabolism, biosynthesis of cuticular hydrocarbon and pheromone ([14, 20], reviewed in [21, 22]). Oenocytes coordinate with fat body in mobilizing lipid storage upon nutrient deprivation [14, 23, 24]. Recent studies in the yellow fever mosquito *Aedes aegypti* showed that pupal oenocytes highly express cytochrome P450 genes, suggesting an important role of oenocytes in detoxification [25]. Despite its roles in lipid metabolism and wax production, we know very little about oenocyte’s other physiological functions, including its role in the regulation of aging and longevity. It is known that aging oenocytes undergo dramatic morphological changes (e.g., increases in cell size and pigmented granules [26]) and exhibit dysregulation of mitochondrial chaperone *Hsp22* [27]. However, transcriptional characterization of oenocyte aging has not been previously performed.

Here, we utilized RiboTag technique [28] to profile the genome-wide changes in ribosome-associated transcripts during oenocyte aging in *Drosophila*. We show that aging and paraquat (PQ) exhibit common and distinct regulation on adult oenocyte translatome. Gene ontology and gene set enrichment analysis (GSEA) revealed that ribosome, proteasome, peroxisome, xenobiotic metabolism, fatty acid metabolism, and DNA replication pathways were altered under aging and oxidative stress. Comparing tissue-specific transcriptomes further revealed that oenocytes were enriched with genes involved liver-like functions (e.g., ketogenesis). Aging oenocytes also shared many molecular signatures with aging liver. Taken together, our translatome analysis revealed a conserved molecular mechanism underlying oenocyte and liver aging. Our study will offer new opportunities for future dissection of novel roles of oenocytes in lipid metabolism, stress response, and aging control.

## Results

### Characterization of age-related changes in ROS production in *Drosophila* oenocytes

In *Drosophila*, larval and adult oenocytes exhibit distinct morphological characteristics [21]. Larval oenocytes are clustering along the lateral body wall [14], while adult oenocytes (used in the present study) appear as segmental dorsal stripes and ventral clusters nearby the abdominal cuticle (Fig. 1A). As oxidative stress is commonly observed in aging tissue, we first examined the age-related changes in ROS production in adult oenocytes. As shown in Figs. 1B&1C, both aging and PQ (an oxidative stress inducer) significantly increased ROS levels in adult oenocytes. Increases in cell and nuclear sizes were also seen in aged oenocytes (Figs. 1B, S1). In the present study, oenocytes were dissected from two ages, 10 days (young) and 30 days (middle age). Middle age was used because many epigenetic and transcriptional changes have been previously observed in the midlife [29-31]. Since elevated ROS levels were already apparent at middle age, a comparison between young and middle age will allow us to capture the early-onset age-related changes in adult oenocytes. Additionally, we noticed that young oenocytes showed much higher induction of ROS under PQ treatment than the oenocytes from middle age (Fig. 1C), suggesting the response to oxidative stress was altered in aged oenocytes.

**Figure 1.**
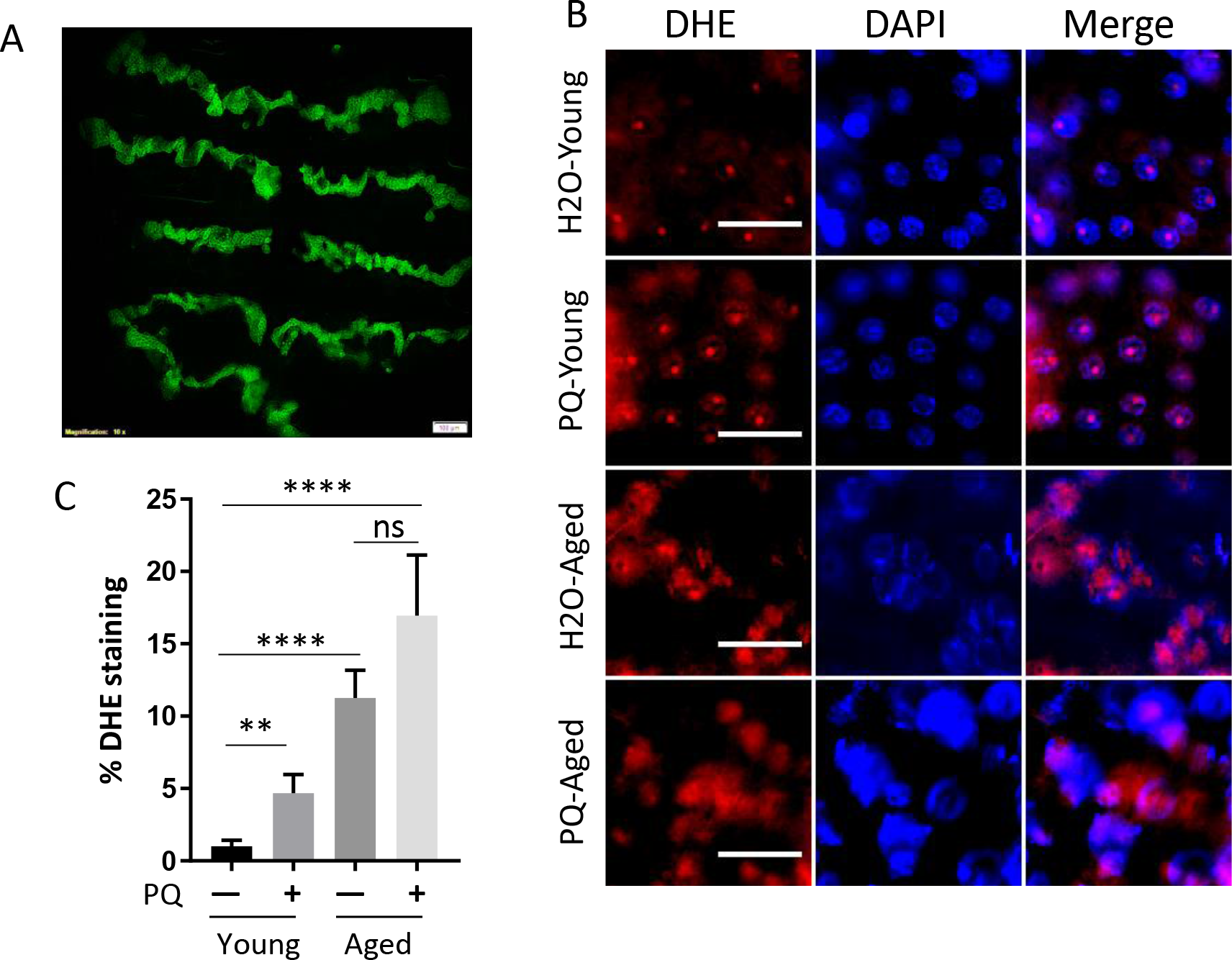
Characterization of age-related changes in ROS production in *Drosophila* oenocytes. (A) Fluorescent image of GFP-labeled oenocytes from the abdomen of *PromE-Gal4; UAS-CD8::GFP* female flies. Scale bar: 100 μm. (B) ROS levels indicated by DHE staining in female oenocytes under aging and paraquat (PQ) treatment. Young: 10-day-old, Aged: 30-day-old. DAPI stains for nuclei. Scale bar: 10 μm. (C) Quantification of DHE staining from Panel (B). One-way ANOVA (*** p<0.001, * p<0.05). N=5.

### Oenocyte-specific translatomic profiling through RiboTag sequencing

Besides their roles in metabolic homeostasis, hydrocarbon and pheromone production (reviewed in [21]), the role of oenocytes in aging regulation has not been carefully examined. Characterization of age-related transcriptional changes in oenocytes is an important step toward our understanding of oenocyte aging. To date, only a few oenocyte transcriptome analyses have been reported [23, 25]. Most of these studies used dissected oenocytes, which often have issues with tissue cross-contamination. To overcome this issue, we performed an oenocyte-specific RiboTag analysis. In the analysis, oenocyte-specific driver *PromE-Gal4* was used to drive the expression of FLAG-tagged *RpL13A*. According to RNA-seq database (at Flybase.org) and a recent ribosomal proteome analysis [32], RpL13A is one of the highly and ubiquitously expressed components in *Drosophila* large ribosomal subunit. Our experimental design facilitates the enrichment of oenocyte-specific ribosome-associated mRNAs and translatomic profiling (Fig. 2A). To verify the efficiency and specificity of our RiboTag profiling, we performed a qRT-PCR analysis to measure the expression of *Desaturase 1 (Desat1)*. *Desat1* is a transmembrane fatty acid desaturase and its E isoform (*desat1-E*) was known to be specifically expressed in female oenocytes [20]. We found that the expression of *desat1-E* was much higher in anti-FLAG immunoprecipitated sample (oenocytes) compared to the input (whole body), suggesting that our RiboTag approach can effectively detect the gene expression from adult oenocytes (Fig. 2B).

**Figure 2.**
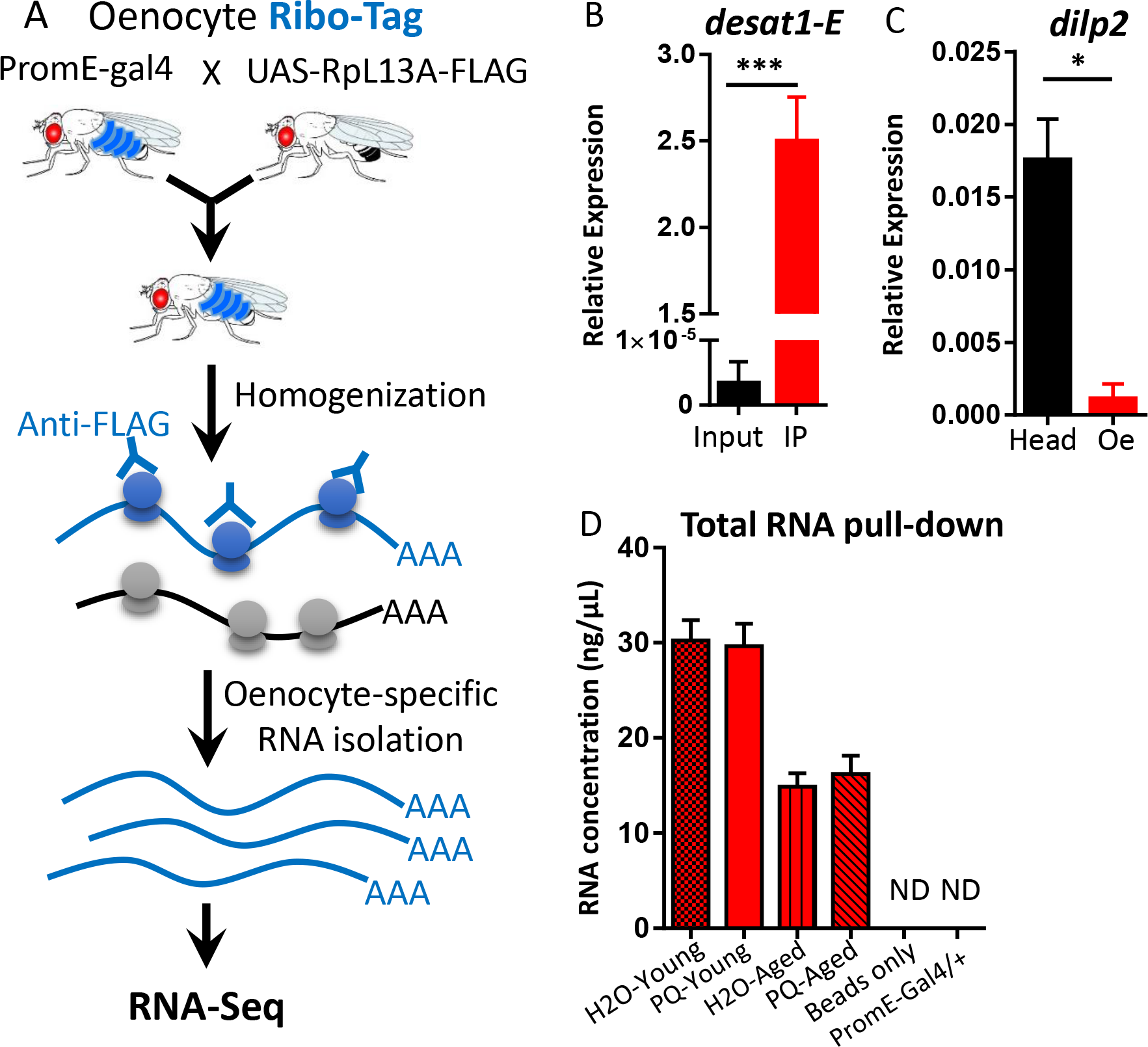
Oenocyte-specific translatomic profiling through RiboTag sequencing. (A) Schematic diagram showing RiboTag procedures. FLAG-tagged ribosomal protein RpL13A was first ectopically expressed in oenocytes. Translating RNAs were immunoprecipitated using antiFLAG antibodies. RNAs were further purified and used in RNA-seq analysis. (B) Oenocyte-specific transcript *desat1-E* highly expressed in anti-FLAG immunoprecipitated sample (IP) compared to the input (whole body lysate). (C) The transcripts of brain-specific gene *Dilp2* was enriched in head samples compared to oenocyte RiboTag samples. One-way ANOVA (**** p<0.0001, *** p<0.001, ** p<0.01, * p<0.05, ns = not significant). N=3. (D) RNA concentrations of various immunoprecipitated samples. ND: Not detected. 200 female flies were used in each condition. Three biological replicates per condition.

To confirm the specificity of the RiboTag analysis, we measured the expression of a brain-specific gene, *insulin-like peptide 2* (*Dilp2*), and found that *Dilp2* expression in oenocyte RiboTag samples was very low compared to the head samples (Fig. 2C). Thus our RiboTag analysis has very little contamination from other tissues (such as brain). We also set up two control experiments to test the specificity of the reagents used in our pull-down assay: 1) Immunoprecipitation of *PromE>RpL13A-FLAG* expressing females using only protein G magnetic beads without adding FLAG antibody. 2) Immunoprecipitation of *PromE-gal4* flies using both Protein G magnetic beads and FLAG antibody. No detectable RNAs were pulled down from the two control groups, suggesting there is none or very little non-specific binding from FLAG antibodies or protein G magnetic beads during the immunoprecipitation (Fig. 2D). Notably, the total RNA pulled down from aged samples were less than those from young oenocytes. This is probably due to age-related decreases in general transcription and translation, because the *PromE-gal4* driver activity remained the same during aging (Fig. S1). Due to the variation in RNA quantity among different samples, we used equal amount of RNAs for all library construction. To examine age- and stress-related transcriptional changes in *Drosophila* oenocytes, we performed RiboTag sequencing on four different experimental groups: H2O-Young, PQ-Young, H2O-Aged, PQ-Aged (see Methods for more details). Female flies were used in the present study, because previous studies showed that *PromE-gal4* drives expression in testis (additional to oenocytes) in male flies [20].

### Differential gene expression (DGE) analysis reveals common and distinct transcriptional regulation by aging and oxidative stress

Using Illumina sequencing (HiSeq 3000, single-end, a read length of 50 base pair), we obtained a total of 402 million reads from 12 library samples (about 11.6X coverage per library). On average, 82.43% of unique reads were mapped to annotated *Drosophila* reference genome. To visualize how gene expression varies under different conditions, we performed principal component analysis (PCA) on the fragments per kilobase million (FPKM) reads. The first component accounts for 50% of the variance and the second component accounts for 9% of variance (Fig. 3A). The PCA analysis showed that three replicates of each condition cluster together, except for one of the H2O-young samples. Two age groups were also well separated. Interestingly, there was a reduced variation between H2O and paraquat treatment in aged oenocytes compared to the young ones (Fig. 3A).

**Figure 3.**
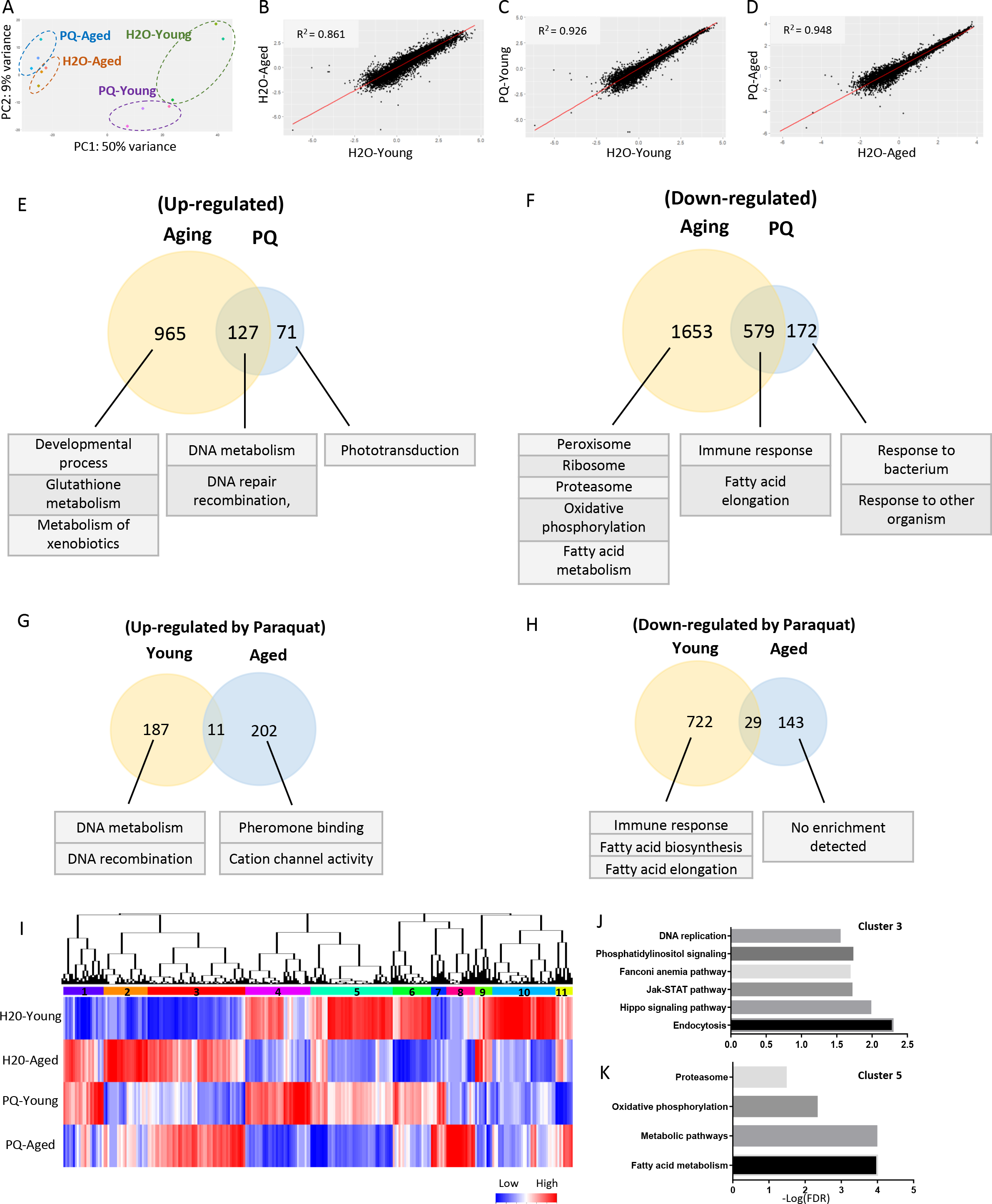
Differential gene expression analysis reveals common and distinct transcriptional regulation by aging and oxidative stress. (A) Principal component analysis (PCA) on four oenocyte translatomes. (B-D) Correlation analysis on the gene expression between H2O-Young and H2O-Aged; H2O-Young and PQ-Young; H2O-Aged and PQ-Aged. Log10 (FPKM) was used in the analysis (E-F) Venn diagram and GO terms for the genes commonly and differentially regulated by aging and paraquat. (G-H) Venn diagram and GO terms for the genes commonly and differentially regulated by paraquat at two ages. (I) Hierarchy clustering analysis on oenocyte translatome. (J-K) Gene ontology analysis on cluster 3 and 5 in panel (I).

DGE analysis was performed using Cufflinks and Cuffdiff tools (fold change ≥ 2, FDR adjusted p-value ≤ 0.05, only protein-coding genes were analyzed). To compare the impacts of aging and oxidative stress on transcriptional changes in adult oenocytes, we first performed correlation analysis using Log-transformed FPKM reads from all four groups. The coefficient of determination (R^2^) was 0.861 between H2O-aged and H2O-young groups (Fig. 3B), 0.926 between H2O-young and PQ-young (Fig. 3C), 0.948 between PQ-aged and H2O-aged (Fig. 3D). Aging induced a bigger transcriptional shift compared to paraquat treatment. Although the change of R^2^ was relatively small, the total number of age-regulated genes was much higher than that under paraquat treatment (Figs. 3E&3F). Thus, both PCA and correlation analyses suggest that aging and paraquat exhibit different impacts on oenocyte translatome.

DGE analysis identified 3324 genes that were differentially expressed during oenocyte aging (1092 up-regulated and 2232 down-regulated), while 949 genes (198 up-regulated and 751 down-regulated) were regulated by paraquat treatment at young ages (Figs. 3E&3F) (Table S1: List 1-4). About 706 DEGs were commonly regulated by aging and paraquat (127 up-regulated and 579 down-regulated) (Table S1: List 5-6). The genes commonly up-regulated by aging and PQ were involved in DNA metabolism, DNA repair and recombination (Fig. 3E), while those commonly down-regulated genes were involved in immune response and fatty acid elongation (Fig. 3F) (Table S2: List 1-4).

Besides common transcriptional regulation between aging and oxidative stress, many genes were differential regulated between the two processes. A total of 2618 genes (965 up-regulated and 1653 down-regulated) were only regulated by aging (Figs. 3E&3F) (Table S1: List 5-6). Genes up-regulated in aged oenocytes were enriched in several Gene ontology (GO) terms, including developmental process, glutathione metabolism and metabolism of xenobiotics (Table S2: List 5). The down-regulated genes are enriched in peroxisome, ribosome, proteasome, oxidative phosphorylation, and fatty acid metabolism (Table S2: List 6). About 243 genes (71 up-regulated and 172 down-regulated) were only regulated by paraquat treatment at young ages. These genes are enriched for biological processes like response to bacterium, response to other organism, and phototransduction (Table S2: List 7-8).

It is known that stress tolerance declines with age [33], which can be caused by impaired transcriptional regulation of stress signaling pathways [34]. Our transcriptome analysis showed that the total number of PQ-regulated genes decreased with aging (Figs. 3G&3H). About 949 genes were differentially expressed under paraquat treatment at young ages (198 up-regulated and 751 down-regulated), while only 385 genes were differentially expressed at middle ages (213 up-regulated and 172 down-regulated) (Table S1: List 7-8). In addition, paraquat treatment targeted a different sets of the biological processes and signaling pathways between young and middle ages (Figs. 3G&3H). In young oenocytes, paraquat up-regulated pathways like response to DNA metabolism and DNA recombination, while down-regulating immune response, fatty acid biosynthesis, and fatty acid elongation (Table S2: List 9-10). In contrast, different sets of pathways were up-regulated by paraquat at middle ages, such as pheromone binding and cation channel activity. No pathway was found enriched for genes down-regulated by paraquat at middle ages (Table S2: List 11-12).

Next, we performed hierarchical clustering analysis and identified 11 distinct clusters among four groups (Fig. 3I). Among 11 clusters, cluster 3 and 5 are two major clusters. Cluster 3 includes genes that were up-regulated in aged oenocytes compared to young ones. Gene ontology analysis showed that cluster 3 was enriched with genes in endocytosis, hippo, JAK-STAT, fanconi anemia pathway, phosphatidylinositol signaling, and DNA replication (Fig. 3J). Cluster 5 consisted of genes down-regulated by aging, and was enriched in fatty acid metabolism, oxidative phosphorylation, and proteasome (Fig. 3K). Taken together, our RiboTag analysis revealed common and distinct transcriptional changes under aging and oxidative stress in adult oenocytes.

### Gene set enrichment analysis (GSEA) reveals up- and down-regulated pathways in aged oenocytes

To further characterize oenocyte-specific signaling pathways that were regulated by aging and oxidative stress, we performed gene set enrichment analysis (GSEA) using a collection of pre-defined gene sets retrieved from Kyoto Encyclopedia of Genes and Genomes (KEGG) database. Through GSEA, we discovered five pathways within which genes were up-regulated with age (FDR q-value<0.05) (Figs. 4A&4C) (Table S2: List 13). They are mismatch repair, DNA replication, base excision repair, nucleotide excision repair, and fanconi anemia pathways. These pathways were tightly related to the cellular responses to DNA replication stress, suggesting a possible increased DNA replication stress during oenocyte aging. Several key players in DNA replication stress response were up-regulated aged oenocytes, such as ATR/mei-41 (ATM- and Rad3-related kinase) and TopBP1/mus101 (DNA topoisomerase 2-binding protein 1).

**Figure 4.**
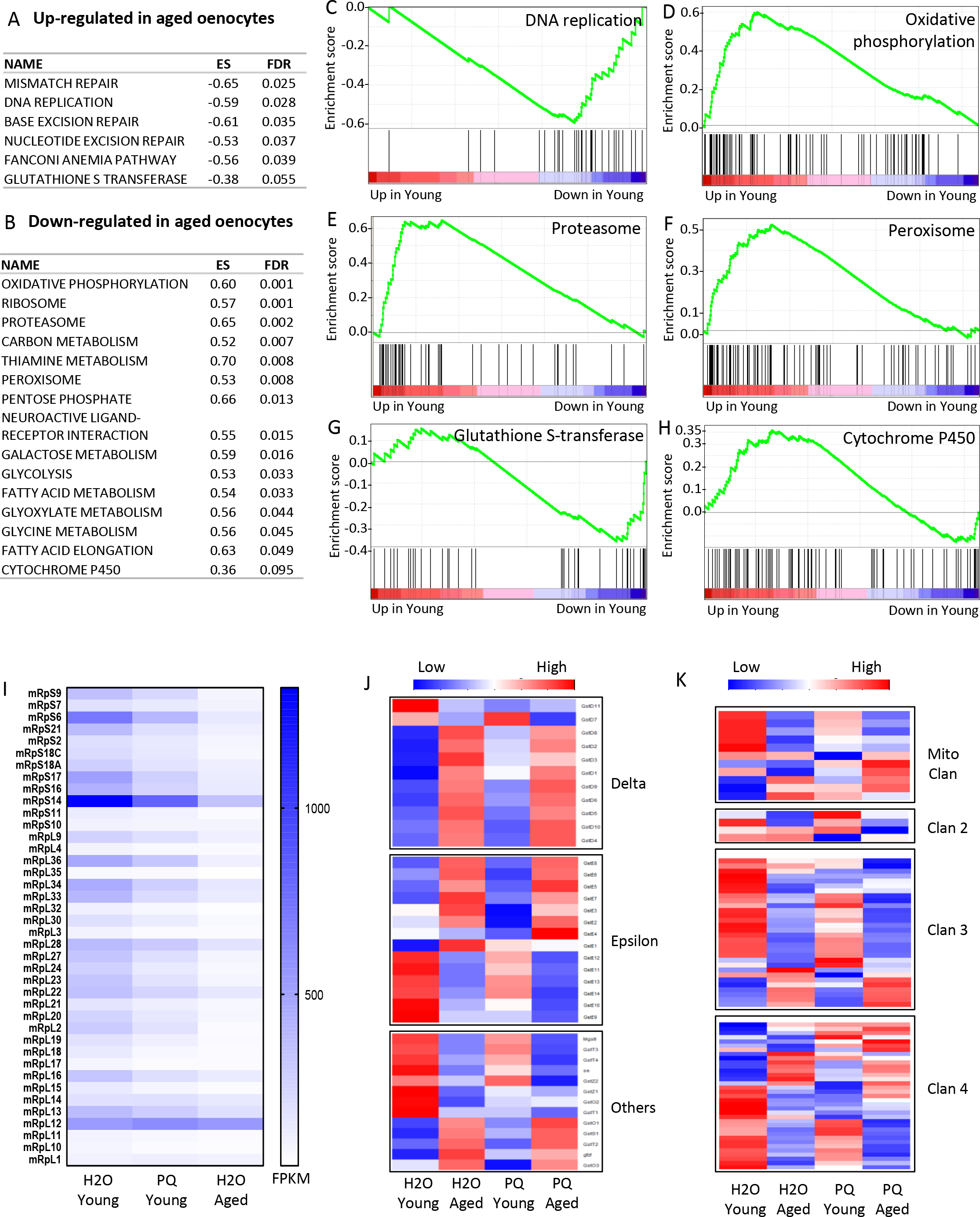
GSEA analysis revealed up- and down-regulated pathways under aging. (A) List of the pathways up-regulated in aged oenocytes. (B) List of the pathways down-regulated in aged oenocytes. ES: Enrichment score. (C-H) GSEA enrichment profiles of six pathways: DNA replication, oxidative phosphorylation, proteasome, peroxisome, glutathione S-transferase, cytochrome P450. (I-K) Heatmaps for mitochondrial ribosomal subunits, glutathione S-transferase, cytochrome P450.

On the other hand, GSEA analysis revealed 14 pathways within which most of genes were significantly down-regulated during aging, such as oxidative phosphorylation, ribosome, proteasome, and peroxisome (Figs. 4B, 4D, 4E, 4F) (Table S2: List 13). These results suggest that the functions of many key cellular organelles/components (e.g., mitochondria and peroxisome) were impaired in aged oenocytes. In aged oenocytes, we found that the key components of all five complexes in mitochondrial electron transport chain were down-regulated, such as NADH dehydrogenase subunits (e.g., *ND-13, ND-15, ND-30, ND-B8*), succinate dehydrogenase (e.g., *SdhC, SdhD*), cytochrome bc1 complex (e.g., *Cyt-c1, UQCR-14, UQCR-C2, UQCR-Q, ox*), cytochrome c oxidase subunits (e.g., *COX4, COX5A, COX5B*), and ATP synthase subunits (e.g., *ATPsynB, ATPsynD, ATPsynF, ATPsynO*) (Table S1: List 2). Interestingly, we found that aging down-regulated many mitochondrial ribosomal subunit genes (44 out of 72 annotated mitochondrial ribosomal proteins) (Fig. 4I) (Table S1: List 2). Lastly, we observed an age-related decrease in the expression proteasome subunit genes. These include 20S protein subunits (e.g., *Prosalpha2, Prosalpha3, Prosbeta1, Prosbeta2, Prosbeta3*), and 19S regulatory cap subunits (e.g., *Rpn1, Rpn11, Rpn12, Rpt1, Rpt2, Rpt3*) (Table S1: List 2). The analysis on peroxisome function is described in a following section.

Reduced xenobiotic metabolism is one of the hallmarks of liver aging [35]. Xenobiotics metabolism (or detoxification) consists of three major phases [36]. The Phase I and II enzymes represent the most abundant classes of detoxification system, including cytochrome P450 (CYPs) and glutathione S-transferases (GSTs). Interestingly, our GSEA analysis revealed distinct expression patterns for these two detoxification enzyme families. We found that almost all of the GSTs in Delta class were up-regulated under aging, while other classes showed mixed expression patterns (Fig. 4J) (Table S1: List 9). The microsomal glutathione S-transferase (*Mgstl*), one of the highly enriched oenocyte genes, was significantly down-regulated during oenocyte aging (Table S1: List 9).

On the other hand, most of the cytochrome P450 genes were down-regulated in aged oenocytes (Figs. 4H&4K). Many of the down-regulated CYPs have been previously linked to insecticide resistance or xenobiotic metabolism, such as *Cyp6a8*, *Cyp6a21, Cyp308a1, Cyp12a4, Cyp6a2, Cyp6w1, and Cyp313a1* (Table S1: List 10). Besides metabolizing exogenous chemicals, several CYPs catalyze endogenous metabolites and play key roles steroid hormone biosynthesis and fatty acid metabolism. For example, *Cyp4g1* is a key CYP gene involved in cuticular hydrocarbon biosynthesis [37] and triglyceride metabolism [14]. The expression of *Cyp4g1* was decreased in aged oenocytes (Table S1: List 10). About eight CYPs (also known as the Halloween genes) in *Drosophila* that are known to regulate ecdysteroid metabolism. Two of them, *Cyp306a1* (*Phantom*) and *Cyp315a1* (*Shadow*), were highly expressed in oenocytes (32-fold and 12.5-fold enriched respectively) (Fig. S2). During oenocyte aging, *Phantom* was down-regulated, whereas *Shadow* was up-regulated (Table S1: List 10).

### Peroxisome pathway is transcriptionally deregulated in aged oenocytes

Recent studies suggest that peroxisome protein import is impaired during aging [38]. Our GSEA analysis revealed that except for *Pex1* (up-regulated), most of the genes involved in peroxisome biogenesis (also called peroxin, *PEX*) were down-regulated in aged oenocytes (Figs. 4F&5A) (Table S1: List 11). Out of 16 peroxin genes, five of them showed significant down-regulation during aging (fold change ≥ 2, FDR adjusted p-value ≤ 0.05). They are matrix enzyme import components (*Pex5, Pex12*), receptor recycling (*Pex6*), and membrane assembly components (*Pex16, Pex19*) (Figs. 5A&5B). In addition, most of the *PEX* genes were also down-regulated by paraquat treatment, but to a less extend comparing to aging (Fig. 5C).

**Figure 5.**
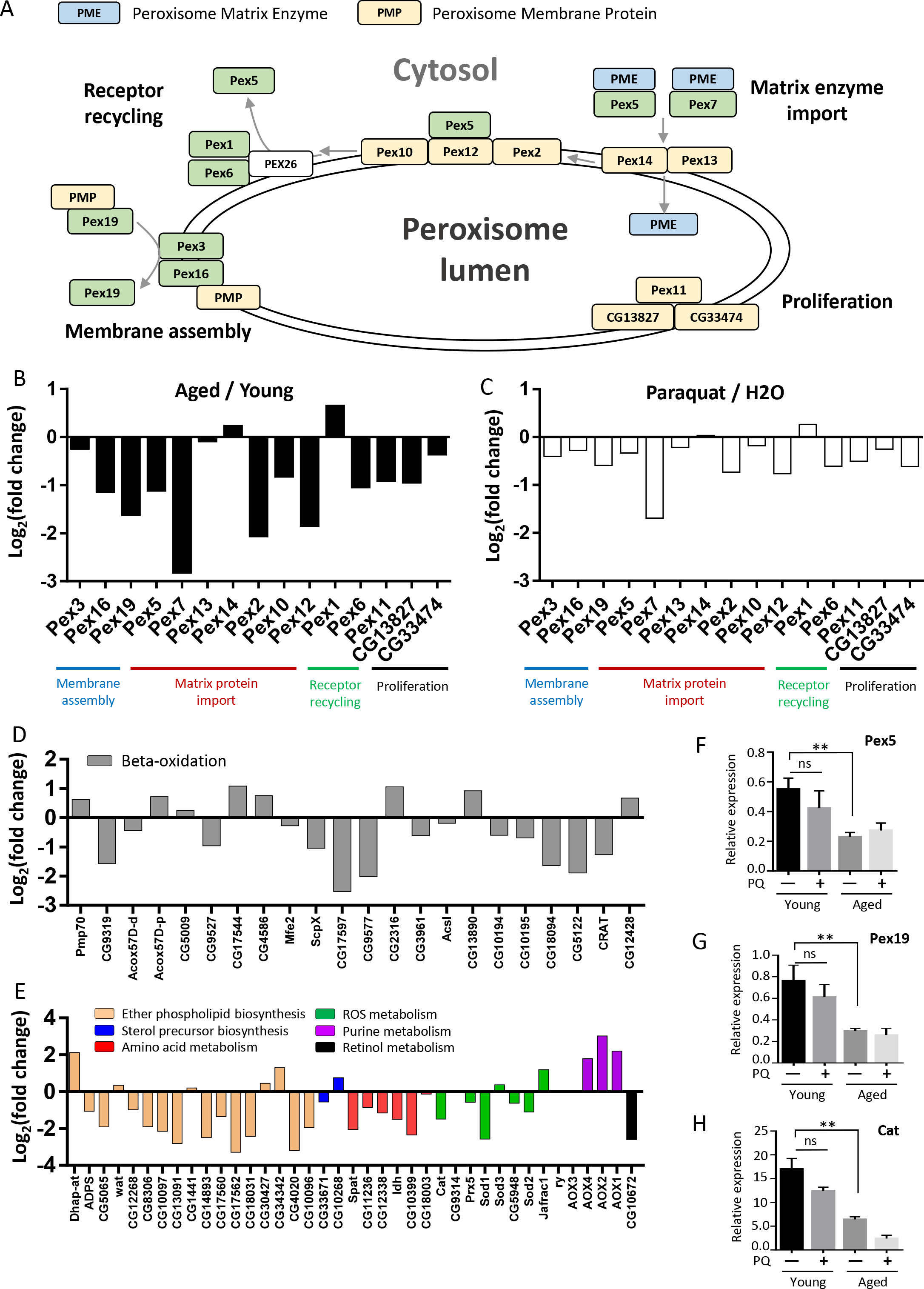
Peroxisome pathway is transcriptionally deregulated in aged oenocytes. (A) Schematic diagram showing peroxisome pathway and the role of each peroxin (PEX) genes. (B-C) Log_2_ (fold change) of the expression of PEX genes under aging and paraquat treatment, based on oenocyte RiboTag sequencing results. (D-E) Log2 (fold change) of the expression of genes involved in other peroxisome functions during oenocyte aging. (F-H) qRT-PCR verification of three peroxisome genes (*Pex5, Pex19, Cat*). One-way ANOVA (, ** p<0.01, ns = not significant). N=3.

Besides peroxisome biogenesis, genes involved in other peroxisomal functions were also down-regulated during oenocyte aging (Figs. 5D&5E) (Table S1: List 11). These functions include fatty acid beta-oxidation, ether phospholipid biosynthesis, amino acid metabolism, ROS metabolism, purine metabolism, and retinol metabolism. Several beta-oxidation genes showed significantly decreased expression, including sterol carrier protein X-related thiolases (*ScpX and CG17597*), enoyl-CoA hydratase (*ECH/CG9577*), carnitine O-acetyl-transferases (*CRAT and CG5122*), and nudix hydrolases (*CG10194, CG10195, CG18094*) (Fig. 5D). Consistently, hepatocyte nuclear factor 4 (*HNF4*), the major regulator for mitochondrial and peroxisomal beta-oxidation, was significantly down-regulated under aging and paraquat (Table S1: List 2). On the other hand, a few other beta-oxidative genes were up-regulated in aged oenocyte, such as ABC transporters (*Pmp70, CG2316*) that are responsible for transporting long-chain fatty acids into peroxisome, delta3-delta2-enoyl-CoA isomerase (*PECI/CG13890*), carnitine O-octanoyltransferase (*CROT/CG12428*). Acyl-CoA oxidases (*Acox*) that are involved in the first step of beta-oxidation showed mixed expression pattern (Fig. 5D).

Consistent with increased ROS production during oenocyte aging, most of the genes regulating peroxisomal ROS metabolism were down-regulated in aged oenocytes, such as *catalase* (*Cat*), *superoxide dismutase 1* (*SOD1*), *peroxiredoxin 5* (*Prx5*). Although the majority of ether phospholipid synthesis genes (e.g., fatty acyl-CoA reductase, *FAR*) were down-regulated, there are a few genes that showed up-regulation during aging, such as dihydroxyacetone phosphate acyltransferase (*DHAPAT* or *Dhap-at*), the key enzyme for the production of acyl-DHAP (the obligate precursor of ether lipids). Additionally, three aldehyde oxidases (*Aox1, Aox2, Aox4*) in purine metabolism were up-regulated (Fig. 5E).

To verify our RiboTag sequencing results, we performed qRT-PCR analysis on three selected peroxisome genes, *Pex5, Pex19*, and *Cat*. Consistent with RNA-Seq results, qRT-PCR showed that all three genes were significantly down-regulated in aged oenocytes (Figs. 5F-5H).

### Ketogenesis, fatty acid elongation, and peroxisome pathways are enriched in both oenocytes and liver

Fat body, but not oenocytes, is a long-established tissue model to study liver-and adipose-like functions in *Drosophila* [39]. Although hepatocyte-like functions (e.g., steatosis) have been previously observed in oenocytes [14], it remains unclear whether fat body and oenocytes each perform different aspects of liver-like functions in *Drosophila*. To address this question, we performed a transcriptome comparison among *Drosophila* oenocytes, fat body, and human liver (Table 1).

**Table 1.** Comparison of the GO terms enriched in oenocyte, fat body and liver.

| GO | Oenocyte | Fat body | Liver |
| --- | --- | --- | --- |
| **Biological Process** | Fatty acid biosynthesis<br>Fatty acid elongation<br>Oxidation reduction<br>Immune response<br>Proteasome-mediated protein catabolic process | Carboxylic acid metabolism<br>Immune response<br>Amino acid metabolism | Lipid metabolism<br>Long-chain fatty acid metabolism<br>Oxidation reduction<br>Immune response<br>Xenobiotic metabolism<br>Bile acid metabolism |
| **Molecular Function** | Oxidoreductase activity<br>Threonine-type endopeptidase activity<br>Fatty acid synthase activity<br>Fatty acid elongase activity<br>Peptidoglycan binding | Oxidoreductase activity<br>Metalloendopeptidase activity<br>Heme binding | Oxidoreductase activity<br>Serine-type peptidase activity<br>Lipid binding<br>Heme binding |
| **Cellular Component** | Peroxisome<br>Proteasome | Extracellular region | Peroxisome<br>Extracellular region<br>Endoplasmic reticulum |
| **KEGG Pathway** | Metabolism of xenobiotics<br>Peroxisome<br>Proteasome<br>Ketone body metabolism<br>Biosynthesis of unsaturated fatty acids | Glycine, serine and threonine metabolism<br>Arginine and proline metabolism | Metabolism of xenobiotics<br>Peroxisome<br>Bile secretion<br>PPAR signaling pathway<br>Retinol metabolism<br>Biosynthesis of amino acids |
| **Protein Domain** | ELO family<br>Proteasome subunit | Peptidase S1, M13<br>EGF-like domain | Serpin family<br>Cytochrome P450 |

We first identified genes that were enriched in adult oenocytes by comparing our oenocyte RiboTag data (H2O-Young group) with previously published whole body transcriptome data (Table S1: List 12). Fat body-enriched genes were identified based on *Drosophila* tissue transcriptome database, FlyAtlas [40, 41] (Table S1: List 13). The genes with more than 5-fold higher expression in oenocytes (or fat body) comparing to whole body are defined as oenocyte-enriched (or fat body-enriched) genes. A total of 423 oenocyte-enriched genes and 544 fat body-enriched genes were identified through tissue transcriptome comparison (Fig. S2). A recent study showed that *Drosophila* oenocytes express many liver-like lipid metabolic genes/pathways [14]. About 15 of these genes were also found enriched in our oenocyte translatome analysis (e.g., *Cpr, Cat, spidey, FarO*) (Table S1: List 16). About 463 human liver-enriched genes were retrieved from the Human Protein Atlas [42] (Table S1: List 15).

Interestingly, there was very little overlap between oenocyte-enriched and fat body-enriched genes, suggesting that adult fat body and oenocytes may regulate distinct biological processes (Fig. S2A) (Table S1: List 14). Gene ontology analysis revealed that fat body mainly expressed genes in carboxylic acid and amino acid metabolism, whereas oenocytes were enriched with genes in pathways like fatty acid biosynthesis, fatty acid elongation, proteasome-mediated protein catabolism, xenobiotic metabolism, ketone body metabolism, and peroxisome (Table 1) (Table S2: List 14-15). Furthermore, we found that two innate immunity pathways, Toll and Imd (Immune deficiency), were differentially enriched in fat body and oenocytes (Fig. S2B) (Table S1: List 12-13). Several genes in Imd pathway (*PRGP-LC, PRGP-LB, Dredd*) were enriched in oenocytes, whereas fat body were enriched with genes in Toll pathway (*Tl, PGRP-SA, GNBP3, modSP*) (Fig. S2B). Additionally, most of the antimicrobial peptides (AMPs) were enriched in oenocytes, but not in fat body (Fig. S2B)

When comparing oenocyte and liver transcriptomes, we found that several pathways were specifically enriched in both liver and oenocytes, such as long-chain fatty acid metabolism, peroxisome, and xenobiotic metabolism (Table 1, Table S2: List 14-16). A close look at the enriched genes shared between oenocytes and liver revealed that *HMG-CoA synthase* (*Hmgs* in fly and *HMGCS1/2* in human), the key enzyme involved in ketogenesis and production of β-hydroxy-β-methylglutaryl-CoA (HMG-CoA), was highly expressed in both oenocytes and liver (Figs. 6A&6B). Additionally, two other ketogenesis genes were also highly expressed in both oenocytes and liver. They are *HMG-CoA lyase* (*CG10399* in fly and *HMGCL* in human) and *D-β-hydroxybutyrate dehydrogenase* (*shroud* in fly and *BDH1* in human) (Figs. 6A&6B). Ketogenesis is primarily activated in mammalian liver, especially during fasting. These results suggest that oenocyte may be the fly tissue regulating ketogenesis similar to mammalian liver.

**Figure 6.**
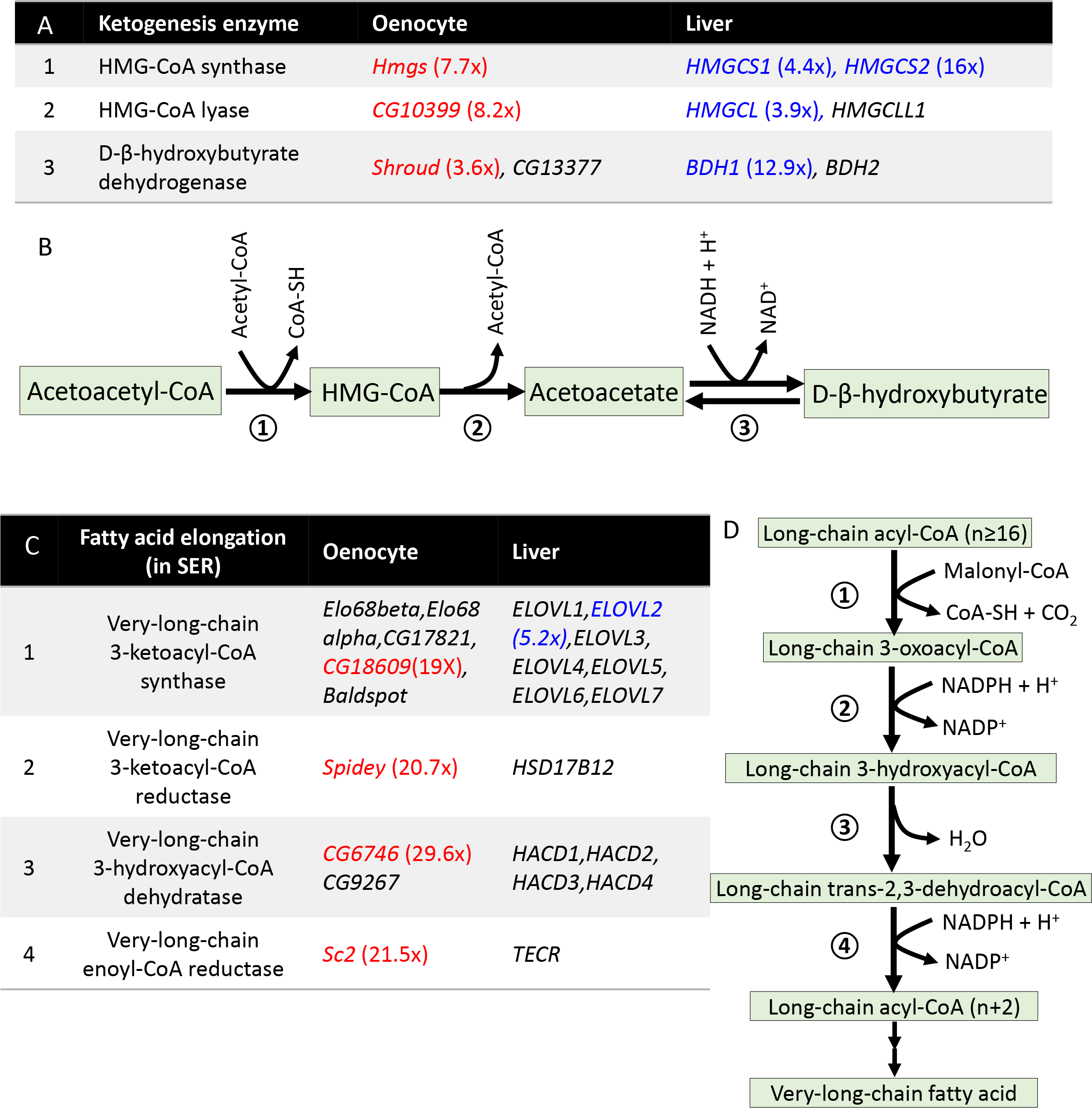
Ketogenesis and fatty acid elongation are enriched in both oenocytes and liver. (A) List of ketogenesis genes that are enriched in both oenocytes and liver. (B) Schematic diagram showing ketogenesis pathway. (C) List of genes in microsomal fatty acid elongation pathway that are enriched in both oenocytes and liver. (D) Schematic diagram showing microsomal fatty acid elongation pathway (in smooth ER). Oenocyte-enriched genes are highlighted in red. Liver-enriched genes are highlighted in blue.

Microsomal fatty acid elongation and the synthesis of very-long-chain fatty acid (VLCFA) were also enriched in both oenocytes and liver (Figs. 6C&6D). Liver and oenocytes were enriched for very-long-chain 3-ketoacyl-CoA synthase (*ELOVL2* in human and *CG18609* in fly), which catalyzes the first step of VLCFA synthesis in smooth endoplasmic reticulum (smooth ER). Oenocytes also showed high expression of three other key enzymes in this process (*spidey, CG6746, Sc2*) (Figs. 6C&6D). The enrichment of fatty acid elongation factors in oenocytes aligns well with previously known oenocyte function in the biosynthesis of VLCFA and hydrocarbons [20, 24]. Notably, several key players involved in the production of cuticular hydrocarbons were enriched in adult oenocytes, including cytochrome P450 *Cyp4g1* (3.4-fold) and its obligatory redox partner, cytochrome P450 reductase *Cpr* (5.7-fold), as well as five peroxisome-localized fatty acyl-CoA reductases (FAR) (*FarO, CG13091, CG14893, CG17562, and CG4020*) (Table S1: List 11-12). In particular, *FarO* was 123-fold enriched in oenocytes, while *CG13091* was 243-fold enriched (Fig. S4).

Additionally, many oenocyte- and liver-enriched genes belong to peroxisome pathway, especially peroxisomal beta-oxidation (*CG17597, CG9577* in oenocytes, *ACOX2, BAAT, EHHADH, ACAA1, SLC27A2, ACSL1, PECR* in liver) (Fig. S4). Genes involved in ROS metabolism (e.g., *Cat, Sod1*) were also enriched in both oenocytes and liver (Fig. S4). Lastly, we found that fibroblast growth factor 21 (*bnl* in fly and *FGF21* in human), a key hormonal factor that regulates glucose homeostasis, was enriched in both oenocytes and liver (Table S1: List 12&15). Taken together, our translatome analysis suggests that oenocytes and fat body regulate distinct processes, and oenocytes may participate several liver-like functions (e.g., ketogenesis, and long-chain fatty acid metabolism).

### Conservation in age-related transcriptional changes between oenocytes and liver

Since our analyses suggest that *Drosophila* oenocytes may perform liver-like functions, we wonder if oenocyte and liver exhibit similar transcriptional changes during aging. To test this, we compared age-related transcriptomic profiles between *Drosophila* oenocytes and mouse liver [10]. We first searched for fly orthologues of mouse liver genes using *Drosophila* Integrative Ortholog Prediction Tool (DIOPT) [43]. Out of 1052 protein-coding genes that are differentially expressed in aging mouse liver, 735 of them have putative orthologues in *Drosophila* genome, corresponding to 881 *Drosophila* genes (Fig. 7A). About 30% of these *Drosophila* orthologues (252 out of 881) also showed differential expression during oenocyte aging, suggesting a large conservation between liver and oenocyte aging (Table S1: List 17-18).

**Figure 7.**
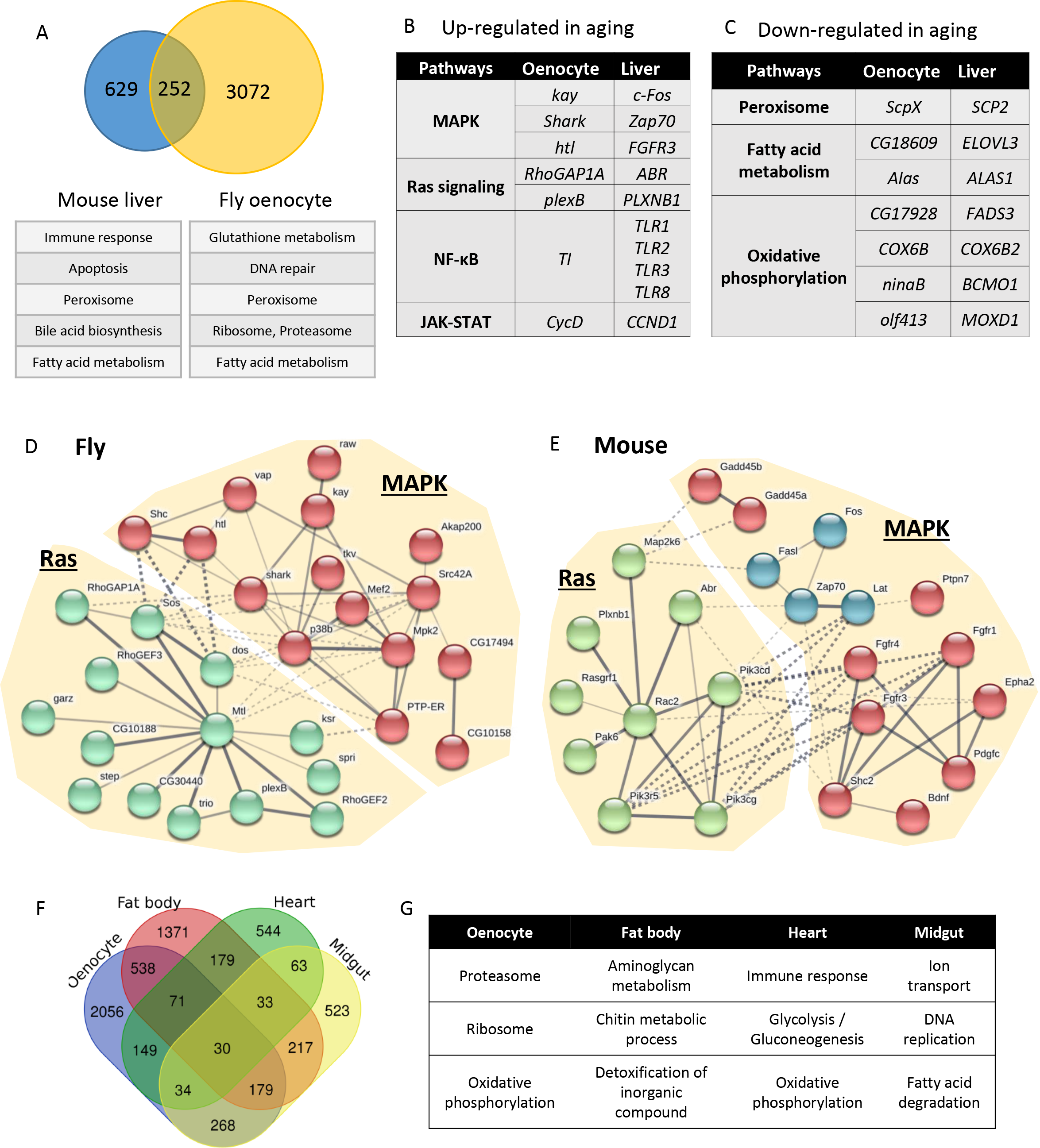
Conservation of age-related transcriptional changes between oenocytes and liver. (A) Venn diagram comparing genes differentially expressed in aged liver and aged oenocytes. The mouse liver genes were first converted to their putative *Drosophila* orthologues before comparing to oenocyte aging genes. GO terms were shown in the lower panel. (B) Signaling pathways that were up-regulated under both oenocyte and liver aging. (C) Signaling pathways that were down-regulated under both oenocyte and liver aging. The genes listed in Panel B&C are the orthologues between *Drosophila* and mouse. (D-E) List of all genes in Ras/MAPK signaling pathway that were down-regulated in aged fly oenocytes and mouse liver. Protein network was generated using STRING (with *kmeans clustering* option). (F) Venn diagram showing the overlap of differentially expressed genes in aged oenocytes, fat body, heart, and midgut. (G) GO terms enriched in aged oenocytes, fat body, heart, and midgut.

Gene ontology analysis showed that several key biological processes were altered in aged liver, including immune response, apoptosis, peroxisome, bile acid biosynthesis, and fatty acid metabolism (Table S2: List 17-18). Among these biological processes, peroxisome and fatty acid metabolism are shared between liver and oenocyte aging (Fig. 7A). Next, we took a close look at the pathways that contain same orthologues between fly and mouse. Genes up-regulated in both aged oenocytes and liver were enriched in pathways like Mitogen-activated protein kinase (MAPK), Ras signaling, NF-κB, and JAK-STAT (Fig. 7B), while down-regulated genes were found in peroxisome, fatty acid metabolism, and oxidative phosphorylation pathways (Fig. 7C) (Table S2: List 17-18). Using STRING protein network analysis, we found that large number of Ras/MAPK signaling components were up-regulated under both oenocyte and liver aging (Figs. 7D&7E), suggesting that age-dependent dysregulation of these pathways are conserved between fly and mammal.

Lastly, we examined age-related transcriptomic changes between oenocytes and several other fly tissues, such as fat body, midgut, and heart. The age-related transcriptional profiles in these fly tissues were obtained from recent genomic studies [44-46] (Table S1: List 19-20). Pathway analysis (using STRING) on these tissue transcriptomes revealed a tissue-specific transcriptional profiles during fly aging (Fig. 7F). Each tissue has its own and unique age-regulated biological processes and pathways (Fig. 7G) (Table S2: List 20-21). For example, genes that were differentially expressed in aged oenocytes are enriched for proteasome and ribosome-related functions, while aged fat body showed transcriptional changes in aminoglycan metabolism, chitin metabolism, and detoxification. In aging heart, immune response, glycolysis and gluconeogenesis were enriched. And ion transport, DNA replication, and fatty acid degradation were altered in aging midgut (Fig. 7G). Taken together, aged oenocytes share similar transcriptional profiles with aging liver, while they also exhibit unique features compared to other fly tissues.

## Discussion

Oenocytes are one of the poorly studied yet important cells in insects [21, 22]. Although previous studies show that oenocytes play a crucial role in lipid metabolism (e.g., synthesis of cuticular hydrocarbon and pheromone), many other oenocyte-regulated physiological functions remain to be determined. Among the uncharacterized functions, we know very little about oenocyte aging and the role of oenocytes in aging regulation. To address these issues, we performed RiboTag sequencing to characterize *Drosophila* oenocyte translatome under aging and oxidative stress. We show that both aging and paraquat up-regulated DNA repair pathway, while down-regulating immune response and fatty acid elongation. In addition, aged oenocytes were associated with impaired peroxisome, mitochondrial, proteasome, and cytochrome P450 pathways. Our RiboTag sequencing also revealed many shared tissue-specific pathways and age-related transcriptional changes between fly oenocytes and mammalian liver, highlighting evolutionarily conserved mechanisms underlying oenocyte and liver aging and potential functional homologies between the two tissues.

### 1. Oenocyte-specific expressed genes are involved in insect-specific and conserved liver-like functions

Previous functional and histological analyses showed that oenocytes contain large amounts of smooth ER and acidophilic cytoplasm (high protein and lipid contents) [47, 48], which is consistent with their roles in lipid synthesis and processing, especially the production of VLCFA and hydrocarbon [20, 24, 49, 50]. Interestingly, *Drosophila* oenocytes uptake and process fatty acids that are released from the storage tissue fat body during food deprivation [14]. The coordination between fat body and oenocytes in mobilizing lipid storage during fasting is quite similar to the adipose-liver axis in mammals. Besides lipid metabolism, many other oenocyte-associated functions (e.g., detoxification and ecdysteroid biosynthesis) have not yet been thoroughly examined at the molecular level. It is unclear whether some of these functions are also conserved liver-like functions, or they are merely insect-specific roles.

To better understand oenocyte function, we conducted oenocyte-specific translatome profiling in adult *Drosophila* and identified 423 genes that were highly expressed in oenocytes (at least 5-fold higher than whole body expression). These genes were enriched in pathways like fatty acid elongation, proteasome-mediated protein catabolism, xenobiotic metabolism, ketogenesis, and peroxisome pathways. There was only a small overlap between oenocyte-enriched and fat body-enriched genes, suggesting that the two tissues regulate distinct functions in *Drosophila*. Comparing to the genes and pathways enriched in human liver, we found that oenocytes shared several biological processes with liver, such as ketogenesis, peroxisomal beta-oxidation, ROS metabolism, long-chain fatty acid metabolism, and xenobiotic metabolism. This is consistent with a previous study showing that *Drosophila* oenocytes expressed high levels of lipid metabolic genes similar to those of mammalian liver [14]. One enriched pathway in *Drosophila* oenocytes that was not observed in the previous study is the ketogenesis pathway. It is well-known that ketone bodies (acetoacetate, β-hydroxybutyrate, and acetone) are primarily produced by liver when glucose is not available as fuel source [51]. Ketogenesis in insects, however, is not well studied. Ketone bodies have been detected in hemolymph, fat body, and thoracic muscle of adult desert locust and cockroach [52-54]. It is speculated that ketone bodies are produced in fat body according to the *ex vivo* tissue culture assay in locust [53]. However, fat body (along with many other tissues) can also oxidize ketone bodies, which is quite different from mammals where the ketogenesis tissue liver cannot oxidize ketone [53]. It might be possible that in previous *ex vivo* tissue culture studies, the ketone production came from a contaminated tissue (like oenocytes), rather than fat body. Based on our oenocyte translatome analysis, most of the ketogenesis genes are highly expressed in oenocytes, but not in fat body. Our data suggest that oenocytes are likely the major ketogenesis tissue. A careful function and genetic analysis, such as cell ablation or tissue-specific gene silencing, will need to be performed to examine whether oenocytes are responsible for ketogenesis in *Drosophila* and in other insect species.

Insect hydrocarbons serve as important waterproofing components, and species- and sex-specific recognition signals. The biosynthesis of hydrocarbons are involved in fatty acid elongation, desaturation, reduction, and oxidative decarbonylation [55]. Our oenocyte translatome analysis revealed an enrichment of genes in microsomal fatty acid elongation system, such as *CG18609, spidey, CG6746*, and *Sc2*. This is consistent with oenocyte’s role in hydrocarbon production and its abundant smooth ER content. In microsomal fatty acid elongation system, *spidey* (also known as *Kar*) encodes for the only very-long-chain 3-ketoacyl-CoA reductase in *Drosophila* genome, and it has been implicated in oenocyte VLCFA synthesis and waterproof of the trachea system [50], as well as the production of cuticular hydrocarbon, ecdysteroid metabolism, and oenocyte maturation [24, 56]. Final two steps of hydrocarbon production in insects are very-long-chain fatty acyl-CoA to aldehydes conversion by FAR and aldehyde oxidative decarbonylation by Cyp4g1 and Cpr [21, 37]. Our translatome analysis showed that five different *FARs* (including *FarO*), *Cyp4g1*, and *Cpr* are highly expressed in adult oenocytes. The large number of FARs expressed in adult oenocytes suggests that aldehyde-forming FARs may be responsible for the production of a variety of hydrocarbons in oenocytes, and each FAR can catalyze a unique set of very-long-chain fatty acyl-CoA esters that vary in saturation status and chain length.

In adult insects (especially in females), ovary is the major tissue for ecdysteroid biosynthesis [57, 58]. It remains to be determined whether other adult tissues are also capable to synthesize ecdysteroids. Interestingly, we found two Halloween genes (*phantom* and *shadow*) that are highly expressed in adult oenocytes, suggesting that oenocytes may participate in ecdysteroid synthesis in adult females. Our findings are consistent with an early study showing that abdominal oenocytes dissected from *Tenebrio molitor* larvae can synthesize 20-Hydroxyecdysone (β-ecdysone) [59]. Several recent studies also detected the expression of Halloween genes in adult tissues other than ovaries, such as brain [60], fat body, muscle, and Malpighian tubule [61]. To functionally verify the role of adult oenocytes in ecdysteroid biosynthesis, direct measurement of ecdysteroid production is needed when Halloween genes are specifically knocked down in oenocytes.

### 2. Impaired peroxisome pathway and fatty acid beta-oxidation are the hallmarks of oenocyte aging

Our translatome analysis identified large number of genes (1092 up-regulated and 2232 down-regulated) that were differentially expressed between young and middle ages, suggesting that dramatic cellular and molecular alterations can be observed in oenocytes at the middle age. Some of these changes are consistent with previous aging transcriptome analysis in *Drosophila* [30, 31, 62], such as up-regulation of DNA repair and down-regulation of oxidative phosphorylation. On the other hand, oenocyte aging was specifically associated with the dysregulation of several other pathways, such as down-regulation of peroxisome and fatty acid metabolism pathways. Peroxisomes are important subcellular organelles that participate in a variety of metabolic pathways, including alpha-oxidation of phytanic acids, beta-oxidation of VLCFA, ether phospholipid synthesis (e.g., plasmalogen biosynthesis), ROS and hydrogen peroxide metabolism, glyoxylate metabolism, catabolism of amino acids and purine [63]. There are about 16 peroxisome biogenesis genes (also known as peroxin, or PEX) in *Drosophila* that are responsible for peroxisome membrane assembly (Pex3, Pex6, Pex9), matrix enzyme import and receptor recycling (Pex5, Pex7, Pex13, Pex14, Pex2, Pex10, Pex12, Pex1, Pex6), and peroxisome proliferation (Pex11) [64]. Mutation in many peroxin genes leads to various forms of peroxisome biogenesis disorder (PBD), also known as Zellweger syndrome (ZS) in human [63]. Our data revealed that aging and oxidative stress decreased the expression of most of the peroxisome biogenesis and protein import genes, which may lead to reduced peroxisome function, including hydrogen peroxide metabolism. Decreased expression of receptor protein Pex5 and reduced peroxisomal enzyme import were previously observed in aged *C. elegans* [38] and during human fibroblast senescence [65]. Among many key peroxisomal enzymes, the importing of antioxidant catalase was significantly affected during fibroblast senescence, which led to accumulation of hydrogen peroxide and further disruption of peroxisome import [65]. Similar to early studies in aging rat liver [66-68], we found that the expression of many peroxisomal antioxidant enzymes (e.g., *Cat, SOD1, Prx5*) decreased in aged oenocytes. The combined dysregulation of peroxisomal gene expression and protein import may attribute to elevated toxic reactive oxygen species, and impaired oenocyte functions. Furthermore, generation of excess peroxisomal ROS could disrupt mitochondria redox balance, leading to mitochondrial dysfunction and tissue aging [69].

Impaired peroxisome biogenesis/protein import during aging not only contributes to reduced antioxidant capacity and elevated ROS levels, but also dysregulation of other peroxisomal functions. Besides ROS metabolism, our translatome analysis revealed that genes involved in peroxisomal beta-oxidation and ether phospholipid were down-regulated under oenocyte aging. This is consistent with previous studies showing that peroxisomal beta-oxidation activity decreased in old mouse liver [70]. Peroxisome has been shown to coordinate with mitochondrial fission/fusion pathway to regulate cellular fatty acid oxidation [71], a major metabolic process dysregulated during mouse aging [72]. Although the metabolic reactions for fatty acid beta-oxidation are similar in mitochondria and peroxisome, a set of fatty acid substrates can only be processed by peroxisomes, such as VLCFA, pristanic acid, di- and trihydroxycholestanoic acid (DHCA and THCA), long-chain dicarboxylic acids, certain polyunsaturated fatty acids [63, 73]. Mutation of peroxisome fatty acid transporter ABCD1 impaired peroxisomal beta-oxidation and caused to accumulation of VLCFAs and neuroinflammation, which is associated X-link neurodegenerative disease adrenoleukodystrophy [74, 75]. Mouse homozygous mutants of ACOX, which catalyzes the first step of peroxisomal beta-oxidation, also showed accumulation of VLCFA and development of microvesicular fatty liver. Although the expression of two *Drosophila* ACOX genes were not significantly altered during oenocyte aging, ScpX (peroxisomal thiolase) was significantly down-regulated. Mice with ScpX mutation showed defects in peroxisome proliferation, hypolipidemia, motor and peripheral neuropathy, as well as impaired catabolism of methyl-branched fatty acids [76]. In addition, reduced peroxisome function can disrupt lipid homeostasis and lipid composition, which could lead to compromised immune response [77, 78].

### 3. Conservation between oenocytes and liver aging

The comparison of aging transcriptomes between fly oenocytes and mouse liver revealed many shared pathways between the two tissues. Among these conserved pathways, MAPK and Ras signaling pathways were significantly up-regulated in both aged oenocytes and liver. MAPK signaling is one of the major regulatory pathways involved in stress responses (e.g., oxidative stress). The typical MAPK pathway includes three branches: c-Jun N-terminal kinase (JNK), p38/MAPK, and extracellular signal-regulated kinase (ERK). Previous studies show that all three MAPK cascades are elevated under aging, probably due to increased oxidative stress [79, 80]. Dysregulated MAPK signaling has been implicated in cancer and neurodegenerative diseases such as Alzheimer’s disease, Parkinson’s disease, and amyotrophic lateral sclerosis (reviewed in [81]). In model organisms (e.g., *Drosophila* and *C. elegans*), activation of JNK and p38/MAPK extended lifespan and improved tissue functions in late life [82-84]. Among many MAPK components identified in our analysis, the activator protein 1 (AP-1) subunit, *Drosophila* Kay and its mouse orthologue c-Fos, were found significantly induced under aging. Both Kay and cFos are basic leucine zipper transcription factors that mediate JNK signaling to regulate cell proliferation, tissue regeneration, stress tolerance [85, 86]. Since JNK signaling is the key regulator for the maintenance of tissue homeostasis in response to intrinsic and extrinsic stresses (e.g., UV irradiation, ROS, DNA damage, inflammatory cytokines, infection), the induction of Kay/c-Fos indicates an up-regulation of JNK signaling and accumulated cellular damages/stress responses in aged oenocytes and liver. In addition, Ras small GTPase pathway, the upstream regulator of MAPK kinase cascades, was also up-regulated during oenocyte and liver aging. The direct role of Ras signaling pathway in longevity regulation has been previously demonstrated in several model organisms [87-90]. Further studies on Ras/MAPK signaling are needed to advance our understanding on the specific contributions of these pathways in oenocyte and liver aging. Nevertheless, the up-regulation of Ras/MAPK signaling pathways can be used as an important molecular signature and biomarker for oenocyte and liver aging.

## Conclusion

Using RiboTag sequencing, we characterized the first oenocyte translatome profiles in *Drosophila*. Our analysis uncovered many previously unexplored oenocyte-specific molecular pathways, especially those associated with oxidative stress and aging. Some of these pathways were found enriched in both fly oenocytes and mammalian liver, suggesting a functional homolog between the two tissues. We believe that the analysis of oenocyte translatome will contribute significantly to our understanding of oenocyte biology, as well as the molecular mechanisms for its role in stress response and aging regulation.

## Methods

### Fly strains, aging and paraquat treatment

Flies are raised in 12h:12 h light:dark cycle at 25 °C, 60% relative humidity on agar-based diet with 0.8% cornmeal, 10% sugar, and 2.5% yeast (unless otherwise noted). Fly strains used in the present study include: *w*\*; *PromE-Gal4* (also known as *Desat1-GAL4.E800*) (Bloomington #65405) [20], *PromE-Gal4; UAS-CD8::GFP* (a gift from Alex Gould), *UAS-RpL13A-FLAG* (a gift from Pankaj Kapahi), To age flies, females were collected two days after eclosion, and twenty females per vial were maintained at 25 °C and transferred to fresh food every 2-3 days. Two ages were tested, young (10-day-old) and middle age (30-day-old). For paraquat treatment, flies were fed on fly food containing 10 mM of paraquat (at the food surface) for 24 hours prior to each assay.

### Dihydroethidium (DHE) staining

Young and aged flies were fed on normal food or paraquat (10 mM) for 24 hours prior to the staining with dihydroethidium (Calbiochem, Burlington, MA, USA. Catalog number: 3848326-0). DHE staining was performed as previously described [91]. Briefly, fly abdomen was dissected out (fat body removed) and incubated with 30 μM of DHE in Schneider’s *Drosophila* Medium (ThermoFisher Scientific, Catalog number: 21720-024) for 5 minutes in a dark chamber on an orbital shaker. After additional 5 minutes incubation with 1 μg/mL of Hoechst 33342 (ImmunoChemistry Technologies, Bloomington, MN, USA. Catalog number: 639), fly abdomen was mounted with 50% glycerol in PBS. DHE staining was visualized with Olympus BX51WI upright epifluorescence microscopy.

### Oenocyte RiboTag

Female progeny from the crosses between *PromE-gal4* and *UAS-RpL13A-FLAG* were collected two days after eclosion. Four different experimental groups were tested: 1). 10-day-old females fed on normal food (H2O-Young); 2). 10-day-old females treated with 10 mM of paraquat for 24 hours (PQ-Young); 3). 30-day-old females fed on normal food (H2O-Aged); 4). 30-day-old females treated with 10 mM of paraquat for 24 hours (PQ-Aged). Three biological replicates (200 females per replicate) were performed for each group. Female flies were used in the present study, because *PromE-gal4* drives expression in testis (additional to oenocytes) in male flies [20].

RiboTag was performed following the protocol from McKnight Lab [28]. Briefly, flies were first frozen and ground in nitrogen liquid. The fly powder was then further homogenized in a Dounce tissue grinder containing 5 mL of homogenization buffer (50 mM Tris-HCl, pH 7.4, 100 mM KCl, 12 mM MgCl2, 1 mM DTT, 1% NP-40, 400 units/ml RNAsin RNase inhibitor, 100 μg/ml of cycloheximide, 1 mg/ml heparin, and Protease inhibitors). After centrifuging the homogenate at 10,000 rpm for 10 minutes, the supernatant was first pre-cleaned using SureBeads^™^ Protein G Magnetic Beads (Bio Rad, Hercules, CA, USA. Catalog number: 161-4023), and then incubated with 15 μl of anti-FLAG antibody (Sigma-Aldrich, St. Louis, MO, USA. Catalog number: F1804) for about 19 hours at 4 °C. The antibody/lysate mixture was then incubated with 100 μl of SureBeads for 3 hours at 4 °C. Ribosome-bound RNA was extracted and purified using RNeasy Plus Micro Kit (Qiagen, Hilden, Germany. Catalog number: 74034).

### Transcriptome library construction and high-throughput sequencing (RNA-Seq)

RNA-Seq libraries were constructed using 300 ng of total RNA and NEBNext Ultra Directional RNA Library Prep Kit for Illumina (New England Biolabs (NEB), Ipswich, MA, USA. Catalog number: E7420). Poly(A) mRNA was isolated using NEBNext Oligo d(T)_25_ beads and fragmented into 200 nt in size. After first strand and second strand cDNA synthesis, each cDNA library was ligated with a NEBNext adaptor and barcoded with an adaptor-specific index. Twelve libraries were pooled in equal concentrations, and sequenced using Illumina HiSeq 3000 platform (single-end, 50 bp reads format).

### RNA-Seq data processing and differential expression analysis

The RNA-Seq data processing was performed on Galaxy, an open source, web-based bioinformatics platform (https://usegalaxy.org) [92]. FastQC was first performed to check the sequencing read quality. Then the raw reads were mapped to *D. melanogaster* genome (BDGP Release 6 + ISO1 MT/dm6) using Tophat2 v2.1.0 [93]. Transcripts were reconstructed using Cufflinks v2.2.1 with bias correction. Cuffmerge (http://cole-trapnell-lab.github.io/cufflinks/) was used to merge together 12 Cufflinks assemblies to produce a GTF file for further differential expression analysis with Cuffdiff v2.2.1.3 [94]. After normalization, differentially expressed protein-coding transcripts were obtained using following cut-off values, false discovery rate (FDR) ≤ 0.05 and fold-change ≥ 2. RNA-Seq read files have been deposited to NCBI’s Gene Expression Omnibus (GEO) (Accession # is GSE112146). Non-coding gene and low expressed genes (FPKM<0.01) were excluded from the analysis.

### Principal component analysis (PCA), heatmap and expression correlation plot

PCA graph was generated using plotPCA function of R package DESeq2 [95]. Heatmaps and hierarchy clustering analysis were generated using heatmap.2 function of R package gplots. (https://cran.r-project.org/web/packages/gplots). Expression data was log2 transformed and all reads were added by a pseudo-value 1. The expression correlation plots were plotted using R package ggplot2 (https://cran.r-project.org/web/packages/ggplot2).

### Oenocyte-enriched genes and tissue-specific aging transcriptome analysis

Oenocyte-enriched genes were identified by comparing our oenocyte RiboTag data (H2O Young group) to the whole body transcriptome profiles from previous studies (two wild-type backgrounds: *w^1118^:* GSM2647344, GSM2647345, GSM2647345. *yw:* GSM694258, GSM694259). The sequencing reads with FPKM ≥ 0.01 were normalized by quantile normalization function using preprocessCore package. (https://www.bioconductor.org/packages/release/bioc/html/preprocessCore.html). Oenocyte-enriched genes were defined as those with 5-fold higher FPKM in oenocytes comparing to whole body. Fat body-enriched genes were obtained similarly by comparing the expression values between adult fat body and whole body (data retrieved from FlyAtlas).

The lists of differentially expressed genes in multiple fly tissues were extracted from previous transcriptome analyses, heart [44], posterior midgut [46], fat body [45]. Venn diagram analysis (http://bioinformatics.psb.ugent.be/webtools/Venn/) was performed to identify overlapping genes between different tissues.

### Gene Set Enrichment Analysis (GSEA)

For GSEA analysis, a complete set of 136 KEGG pathways in *Drosophila* were downloaded from KEGG. Text were trimmed and organized using Java script. Quantile normalized FPKM values for each group were used as input for parametric analysis, and organized as suggested by GSEA tutorial site (GSEA, http://software.broadinstitute.org/gsea/doc/GSEAUserGuideFrame.html) [96]. Collapse dataset to gene symbols was set to false. Permutation type was set to gene set; enrichment statistic used as weighted analysis; metric for ranking genes was set to Signal to Noise.

### Gene ontology and pathway analysis

Functional annotation analysis of differentially expressed genes was performed using STRING. GO terms (Biological Process, Molecular Function, Cellular Component), KEGG pathway, INTERPRO Protein Domains and Features, were retrieved from the analysis. To build Ras/MAPK protein network in STRING, “kmeans clustering” option was used and number of clusters was set to 2 or 3.

### Quantitative real-time polymerase chain reaction (qRT-PCR)

qRT-PCR was performed using Quantstudio 3 Real-Time PCR system and SYBR green master mixture (Thermo Fisher Scientific, Waltham, MA, USA Catalog number: A25778). To determine the most stable housekeeping gene, the Ct values for four housekeeping genes were examined in all twelve cDNA samples obtained from different treatments. Using an Excel-based tool, *Bestkeeper* [97], we confirmed that *Gapdh1* is the least-variable housekeeping gene across samples (Table S3). All gene expression levels were normalized to *Gapdh1* by the method of comparative Ct [98]. Mean and standard errors for each gene were obtained from the averages of three biological replicates, with one or two technical repeats. Primer sequences are available in Table S4.

### Statistical analysis

GraphPad Prism (GraphPad Software, La Jolla, CA, USA) was used for statistical analysis. To compare the mean value of treatment groups versus that of control, either student t-test or oneway ANOVA was performed using Dunnett’s test for multiple comparison.

## Acknowledgements

We thank Bloomington *Drosophila* Stock Center, Pankaj Kapahi, Alex Gould, Joel Levine for fly stocks, reagents and protocols. Michael Baker and DNA Facility at ISU for help with RNA-Seq analysis. This work was supported by NIH/NIA R00 AG048016 to HB, AFAR Research Grants for Junior Faculty to HB, and Glenn/AFAR Scholarships for Research in the Biology of Aging to KH.

## Author Contributions Statement

Conceived and designed the experiments: KH HB. Performed the experiments: KH WC HB. Analyzed the data: KH WC FZ HB. Wrote the paper: KC FZ HB. All authors reviewed and approved the manuscript.

## Competing Interests

The authors declare that no competing interest exists.

---

**Table S1. Gene list tables**

**Table S2. GO term tables**

**Table S3. Housekeeping gene analysis using *Bestkeeper***

**Table S4. Primer list**

**Figure S1.**
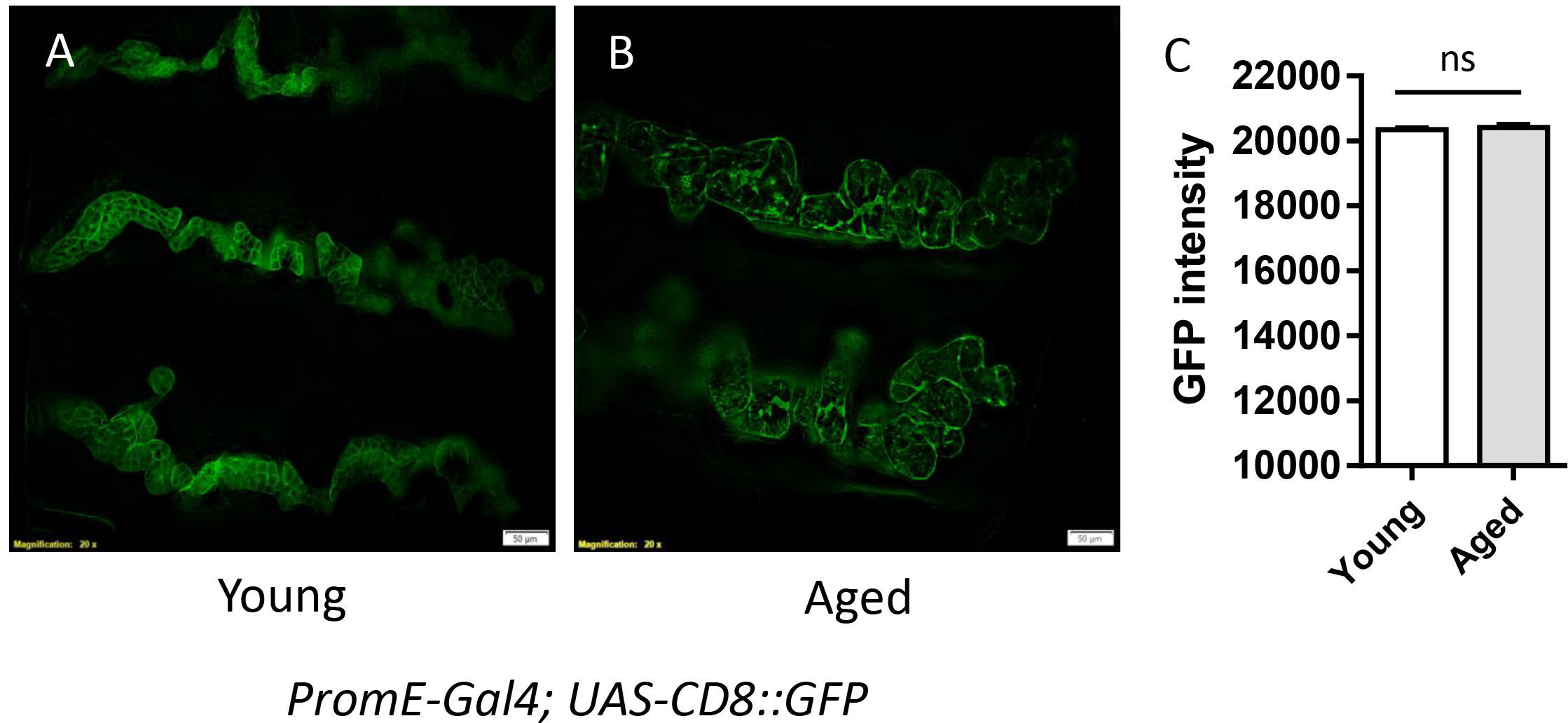
Age-dependent *PromE-gal4* expression pattern. (A-B) Fluorescent image *PromE-Gal4; UAS-CD8::GFP* female flies at two ages: Young (10-day-old), Aged (30-day-old). Scale bar: 50 μm. (C) Quantification of GFP intensity from Panel (A&B). Student t-test (ns = not significant). N=9.

**Figure S2.**
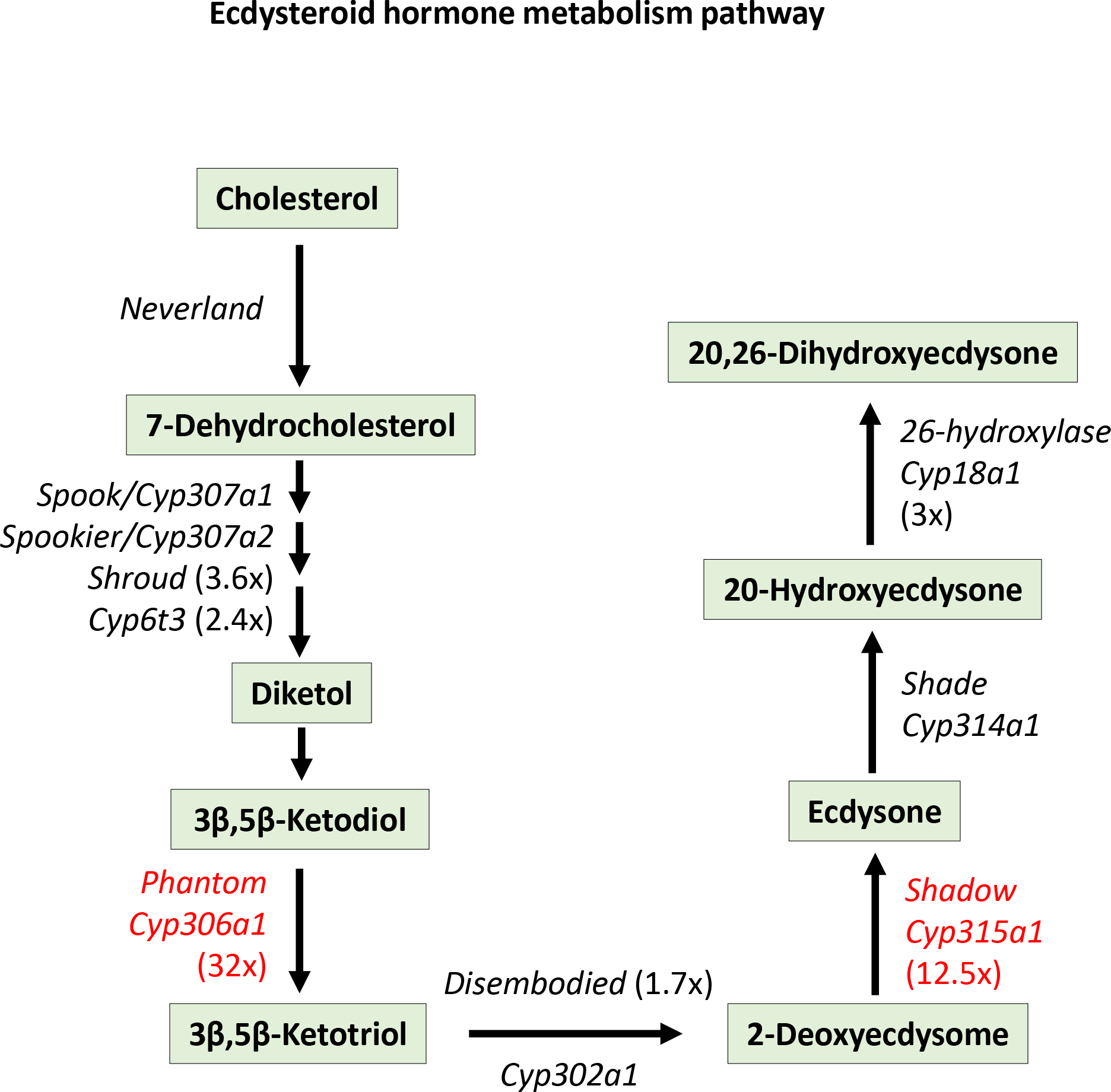
Two ecdysteroid biosynthesis genes highly express in oenocytes. Schematic diagram showing ecdysteroid hormone metabolism pathway. Two Halloween genes, *phantom and shadow*, highly expressed in adult female oenocytes (Highlighted in red).

**Figure S3.**
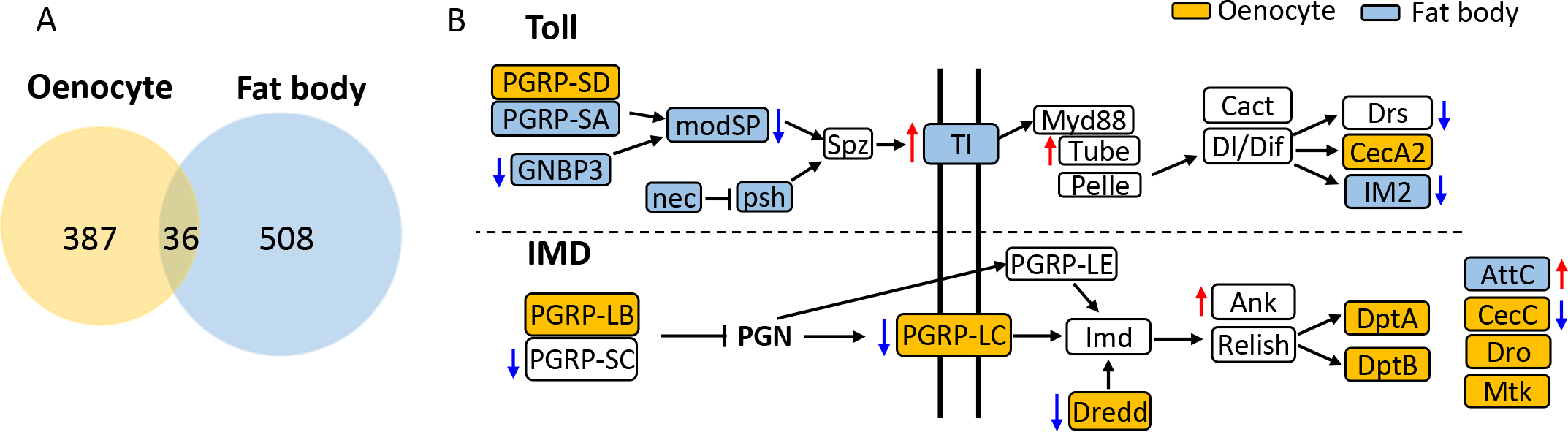
Genes in innate immunity pathway highly express in oenocytes. (A) Genes enriched in oenocytes and fat body show less overlap. (B) Genes in Imd pathway were enriched in oenocytes, while fat body were enriched with genes in Toll pathway (Red arrows denote for age-induced genes. Blue arrows denote for age-repressed gene.).

**Figure S4.**
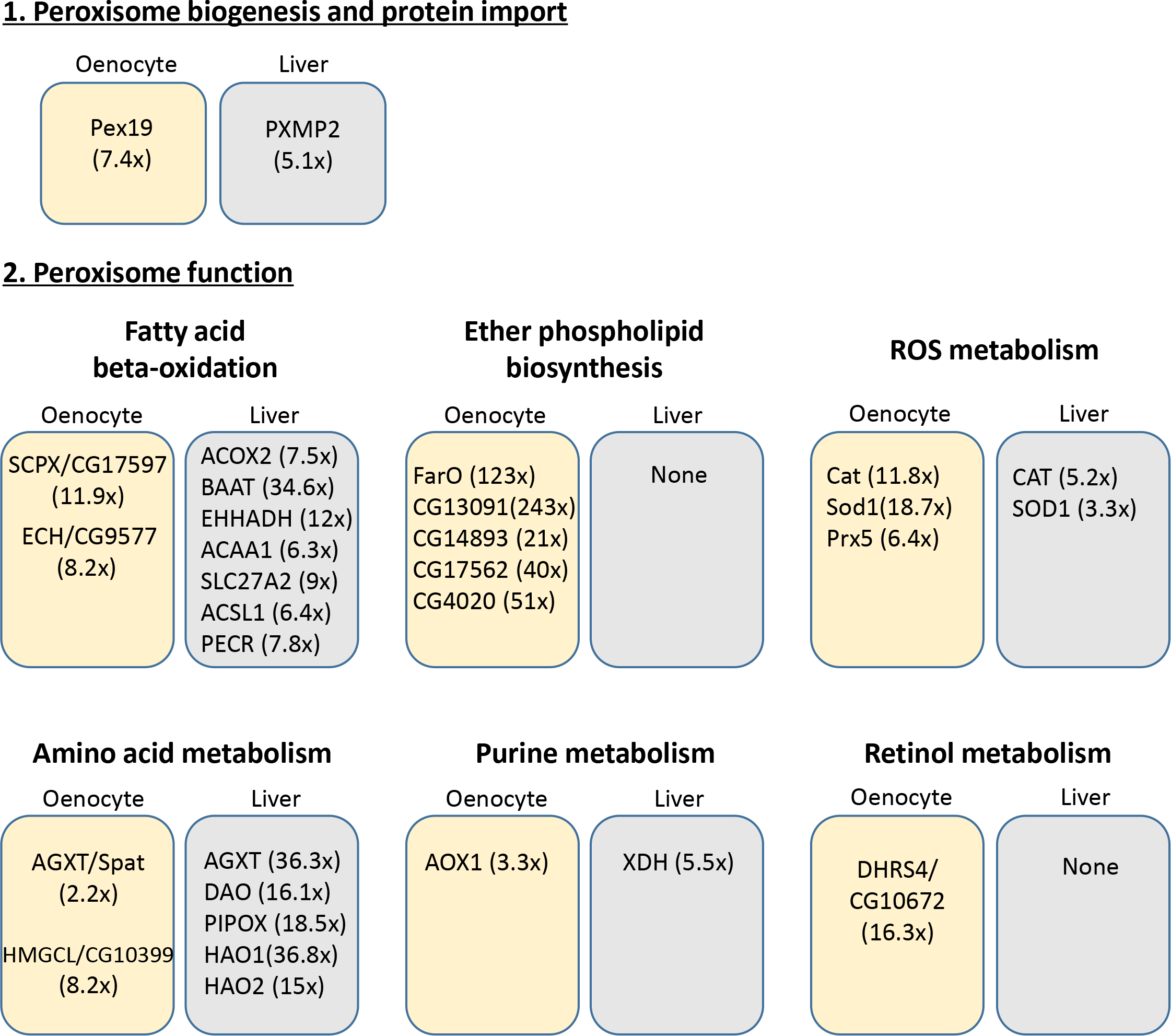
Peroxisome pathways are enriched in both oenocytes and liver. List of peroxisome genes that are enriched in both oenocytes and liver.

## References

1. Kennedy BK, Berger SL, Brunet A, Campisi J, Cuervo AM, Epel ES, Franceschi C, Lithgow GJ, Morimoto RI, Pessin JE et al: Geroscience: linking aging to chronic disease. Cell 2014, 159(4):709–713.

2. Koehler EM, Schouten JN, Hansen BE, van Rooij FJ, Hofman A, Stricker BH, Janssen HL: Prevalence and risk factors of non-alcoholic fatty liver disease in the elderly: results from the Rotterdam study. Journal of hepatology 2012, 57(6):1305–1311.

3. Kim IH, Kisseleva T, Brenner DA: Aging and liver disease. Current opinion in gastroenterology 2015, 31(3):184–191.

4. Sheedfar F, Di Biase S, Koonen D, Vinciguerra M: Liver diseases and aging: friends or foes? Aging Cell 2013, 12(6):950–954.

5. Schmucker DL: Age-related changes in liver structure and function: Implications for disease ? Experimental gerontology 2005, 40(8-9):650–659.

6. Jin J, Iakova P, Breaux M, Sullivan E, Jawanmardi N, Chen D, Jiang Y, Medrano EM, Timchenko NA: Increased expression of enzymes of triglyceride synthesis is essential for the development of hepatic steatosis. Cell Rep 2013, 3(3):831–843.

7. Amir M, Czaja MJ: Autophagy in nonalcoholic steatohepatitis. Expert review of gastroenterology & hepatology 2011, 5(2):159–166.

8. Zhang C, Cuervo AM: Restoration of chaperone-mediated autophagy in aging liver improves cellular maintenance and hepatic function. Nature medicine 2008, 14(9):959–965.

9. Franceschi C, Bonafe M, Valensin S, Olivieri F, De Luca M, Ottaviani E, De Benedictis G: Inflamm-aging. An evolutionary perspective on immunosenescence. Annals of the New York Academy of Sciences 2000, 908:244–254.

10. White RR, Milholland B, MacRae SL, Lin M, Zheng D, Vijg J: Comprehensive transcriptional landscape of aging mouse liver. BMC genomics 2015, 16:899.

11. Tollet-Egnell P, Flores-Morales A, Stahlberg N, Malek RL, Lee N, Norstedt G: Gene expression profile of the aging process in rat liver: normalizing effects of growth hormone replacement. Molecular endocrinology 2001, 15(2):308–318.

12. Horvath S, Erhart W, Brosch M, Ammerpohl O, von Schonfels W, Ahrens M, Heits N, Bell JT, Tsai PC, Spector TD et al: Obesity accelerates epigenetic aging of human liver. Proc Natl Acad Sci U S A 2014, 111(43):15538–15543.

13. He Y, Jasper H: Studying aging in Drosophila. Methods 2014, 68(1):129–133.

14. Gutierrez E, Wiggins D, Fielding B, Gould AP: Specialized hepatocyte-like cells regulate Drosophila lipid metabolism. Nature 2007, 445(7125):275–280.

15. Arrese EL, Soulages JL: Insect fat body: energy, metabolism, and regulation. Annual review of entomology 2010, 55:207–225.

16. Partridge L, Alic N, Bjedov I, Piper MD: Ageing in Drosophila: the role of the insulin/Igf and TOR signalling network. Experimental gerontology 2011, 46(5):376–381.

17. Bai H, Kang P, Tatar M: Drosophila insulin-like peptide-6 (dilp6) expression from fat body extends lifespan and represses secretion of Drosophila insulin-like peptide-2 from the brain. Aging Cell 2012, 11(6):978–985.

18. Hwangbo DS, Gershman B, Tu MP, Palmer M, Tatar M: Drosophila dFOXO controls lifespan and regulates insulin signalling in brain and fat body. Nature 2004, 429(6991):562–566.

19. Giannakou ME, Goss M, Junger MA, Hafen E, Leevers SJ, Partridge L: Long-lived Drosophila with overexpressed dFOXO in adult fat body. Science 2004, 305(5682):361.

20. Billeter JC, Atallah J, Krupp JJ, Millar JG, Levine JD: Specialized cells tag sexual and species identity in Drosophila melanogaster. Nature 2009, 461(7266):987–991.

21. Makki R, Cinnamon E, Gould AP: The development and functions of oenocytes. Annual review of entomology 2014, 59:405–425.

22. Martins GF, Ramalho-Ortigao JM: Oenocytes in insects. Invertebrate Survival Journal 2012, 9:139–152.

23. Chatterjee D, Katewa SD, Qi Y, Jackson SA, Kapahi P, Jasper H: Control of metabolic adaptation to fasting by dILP6-induced insulin signaling in Drosophila oenocytes. Proc Natl Acad Sci U S A 2014, 111(50):17959–17964.

24. Cinnamon E, Makki R, Sawala A, Wickenberg LP, Blomquist GJ, Tittiger C, Paroush Z, Gould AP: Drosophila Spidey/Kar Regulates Oenocyte Growth via PI3-Kinase Signaling. PLoS Genet 2016, 12(8):e1006154.

25. Martins GF, Ramalho-Ortigao JM, Lobo NF, Severson DW, McDowell MA, Pimenta PF: Insights into the transcriptome of oenocytes from Aedes aegypti pupae. Memorias do Instituto Oswaldo Cruz 2011, 106(3):308–315.

26. Johnson MB, Butterworth FM: Maturation and aging of adult fat body and oenocytes in Drosophila as revealed by light microscopic morphometry. Journal of morphology 1985, 184(1):51–59.

27. Tower J, Landis G, Gao R, Luan A, Lee J, Sun Y: Variegated expression of Hsp22 transgenic reporters indicates cell-specific patterns of aging in Drosophila oenocytes. The journals of gerontology Series A, Biological sciences and medical sciences 2014, 69(3):253–259.

28. Sanz E, Yang L, Su T, Morris DR, McKnight GS, Amieux PS: Cell-type-specific isolation of ribosome-associated mRNA from complex tissues. Proc Natl Acad Sci U S A 2009, 106(33):13939–13944.

29. Peleg S, Feller C, Ladurner AG, Imhof A: The Metabolic Impact on Histone Acetylation and Transcription in Ageing. Trends in biochemical sciences 2016, 41(8):700–711.

30. Hall H, Medina P, Cooper DA, Escobedo SE, Rounds J, Brennan KJ, Vincent C, Miura P, Doerge R, Weake VM: Transcriptome profiling of aging Drosophila photoreceptors reveals gene expression trends that correlate with visual senescence. BMC genomics 2017, 18(1):894.

31. Pletcher SD, Macdonald SJ, Marguerie R, Certa U, Stearns SC, Goldstein DB, Partridge L: Genome-wide transcript profiles in aging and calorically restricted Drosophila melanogaster. Curr Biol 2002, 12(9):712–723.

32. Alonso J, Santaren JF: Characterization of the Drosophila melanogaster ribosomal proteome. Journal of proteome research 2006, 5(8):2025–2032.

33. Bloomer SA, Zhang HJ, Brown KE, Kregel KC: Differential regulation of hepatic heme oxygenase-1 protein with aging and heat stress. The journals of gerontology Series A, Biological sciences and medical sciences 2009, 64(4):419–425.

34. Suh JH, Shenvi SV, Dixon BM, Liu H, Jaiswal AK, Liu RM, Hagen TM: Decline in transcriptional activity of Nrf2 causes age-related loss of glutathione synthesis, which is reversible with lipoic acid. Proc Natl Acad Sci U S A 2004, 101(10):3381–3386.

35. Mori K, Blackshear PE, Lobenhofer EK, Parker JS, Orzech DP, Roycroft JH, Walker KL, Johnson KA, Marsh TA, Irwin RD et al: Hepatic transcript levels for genes coding for enzymes associated with xenobiotic metabolism are altered with age. Toxicologic pathology 2007, 35(2):242–251.

36. Xu C, Li CY, Kong AN: Induction of phase I, II and III drug metabolism/transport by xenobiotics. Archives of pharmacal research 2005, 28(3):249–268.

37. Qiu Y, Tittiger C, Wicker-Thomas C, Le Goff G, Young S, Wajnberg E, Fricaux T, Taquet N, Blomquist GJ, Feyereisen R: An insect-specific P450 oxidative decarbonylase for cuticular hydrocarbon biosynthesis. Proc Natl Acad Sci U S A 2012, 109(37):14858–14863.

38. Narayan V, Ly T, Pourkarimi E, Murillo AB, Gartner A, Lamond AI, Kenyon C: Deep Proteome Analysis Identifies Age-Related Processes in C. elegans. Cell systems 2016, 3 (2):144–159.

39. Pandey UB, Nichols CD: Human disease models in Drosophila melanogaster and the role of the fly in therapeutic drug discovery. Pharmacological reviews 2011, 63(2):411–436.

40. Chintapalli VR, Wang J, Dow JA: Using FlyAtlas to identify better Drosophila melanogaster models of human disease. Nat Genet 2007, 39(6):715–720.

41. Leader DP, Krause SA, Pandit A, Davies SA, Dow JAT: FlyAtlas 2: a new version of the Drosophila melanogaster expression atlas with RNA-Seq, miRNA-Seq and sex-specific data. Nucleic Acids Res 2018, 46(D1):D809–D815.

42. Uhlen M, Fagerberg L, Hallstrom BM, Lindskog C, Oksvold P, Mardinoglu A, Sivertsson A, Kampf C, Sjostedt E, Asplund A et al: Proteomics. Tissue-based map of the human proteome. Science 2015, 347(6220):1260419.

43. Hu Y, Flockhart I, Vinayagam A, Bergwitz C, Berger B, Perrimon N, Mohr SE: An integrative approach to ortholog prediction for disease-focused and other functional studies. BMC bioinformatics 2011, 12:357.

44. Monnier V, Iche-Torres M, Rera M, Contremoulins V, Guichard C, Lalevee N, Tricoire H, Perrin L: dJun and Vri/dNFIL3 are major regulators of cardiac aging in Drosophila. PLoS Genet 2012, 8(11):e1003081.

45. Chen H, Zheng X, Zheng Y: Age-associated loss of lamin-B leads to systemic inflammation and gut hyperplasia. Cell 2014, 159(4):829–843.

46. Resnik-Docampo M, Koehler CL, Clark RI, Schinaman JM, Sauer V, Wong DM, Lewis S, D’Alterio C, Walker DW, Jones DL: Tricellular junctions regulate intestinal stem cell behaviour to maintain homeostasis. Nat Cell Biol 2017, 19(1):52–59.

47. Martins GF, Guedes BA, Silva LM, Serrao JE, Fortes-Dias CL, Ramalho-Ortigao JM, Pimenta PF: Isolation, primary culture and morphological characterization of oenocytes from Aedes aegypti pupae. Tissue & cell 2011, 43(2):83–90.

48. Jackson A, Locke M: The formation of plasma membrane reticular systems in the oenocytes of an insect. Tissue & cell 1989, 21(3):463–473.

49. Ferveur JF, Savarit F, O’Kane CJ, Sureau G, Greenspan RJ, Jallon JM: Genetic feminization of pheromones and its behavioral consequences in Drosophila males. Science 1997, 276(5318):1555–1558.

50. Parvy JP, Napal L, Rubin T, Poidevin M, Perrin L, Wicker-Thomas C, Montagne J: Drosophila melanogaster Acetyl-CoA-carboxylase sustains a fatty acid-dependent remote signal to waterproof the respiratory system. PLoS Genet 2012, 8(8):e1002925.

51. Laffel L: Ketone bodies: a review of physiology, pathophysiology and application of monitoring to diabetes. Diabetes/metabolism research and reviews 1999, 15(6):412–426.

52. Bailey EJ, Horne JA, Izatt ME, Hill L: Concentrations of acetoacetate and 3-hydroxybutyrate in pigeon blood and desert locust haemolymph. Life sciences Pt 2: Biochemistry, general and molecular biology 1971, 10(24):1415–1419.

53. Hill L, Izatt MEG, Horne JA, Bailey E: Factors affecting concentrations of acetoacetate and d-3-hydroxybutyrate in haemolymph and tissues of the adult desert locust. Journal of insect physiology 1972, 18(7):1265–1285.

54. Shah J, Bailey E: Enzymes of ketogenesis in the fat body and the thoracic muscle of the adult cockroach. Insect Biochemistry 1976, 6(3):251–254.

55. Howard RW, Blomquist GJ: Ecological, behavioral, and biochemical aspects of insect hydrocarbons. Annual review of entomology 2005, 50:371–393.

56. Chiang YN, Tan KJ, Chung H, Lavrynenko O, Shevchenko A, Yew JY: Steroid Hormone Signaling Is Essential for Pheromone Production and Oenocyte Survival. PLoS Genet 2016, 12(6):e1006126.

57. Gaziova I, Bonnette PC, Henrich VC, Jindra M: Cell-autonomous roles of the ecdysoneless gene in Drosophila development and oogenesis. Development 2004, 131(11):2715–2725.

58. Uryu O, Ameku T, Niwa R: Recent progress in understanding the role of ecdysteroids in adult insects: Germline development and circadian clock in the fruit fly Drosophila melanogaster. Zoological letters 2015, 1:32.

59. Romer F, Emmerich H, Nowock J: Biosynthesis of ecdysones in isolated prothoracic glands and oenocytes of Tenebrio molitor in vitro. Journal of insect physiology 1974, 20(10):1975–1987.

60. Yamazaki Y, Kiuchi M, Takeuchi H, Kubo T: Ecdysteroid biosynthesis in workers of the European honeybee Apis mellifera L. Insect biochemistry and molecular biology 2011, 41(5):283–293.

61. Zheng W, Rus F, Hernandez A, Kang P, Goldman W, Silverman N, Tatar M: Dehydration triggers ecdysone-mediated recognition-protein priming and elevated anti-bacterial immune responses in Drosophila Malpighian tubule renal cells. BMC biology 2018, 16(1):60.

62. Zhan M, Yamaza H, Sun Y, Sinclair J, Li H, Zou S: Temporal and spatial transcriptional profiles of aging in Drosophila melanogaster. Genome research 2007, 17(8):1236–1243.

63. Wanders RJ, Waterham HR: Biochemistry of mammalian peroxisomes revisited. Annual review of biochemistry 2006, 75:295–332.

64. Fujiki Y, Okumoto K, Mukai S, Honsho M, Tamura S: Peroxisome biogenesis in mammalian cells. Frontiers in physiology 2014, 5:307.

65. Legakis JE, Koepke JI, Jedeszko C, Barlaskar F, Terlecky LJ, Edwards HJ, Walton PA, Terlecky SR: Peroxisome senescence in human fibroblasts. Molecular biology of the cell 2002, 13(12):4243–4255.

66. Haining JL, Legan JS: Catalase turnover in rat liver and kidney as a function of age. Experimental gerontology 1973, 8(2):85–91.

67. Semsei I, Rao G, Richardson A: Changes in the expression of superoxide dismutase and catalase as a function of age and dietary restriction. Biochemical and biophysical research communications 1989, 164(2):620–625.

68. Xia E, Rao G, Van Remmen H, Heydari AR, Richardson A: Activities of antioxidant enzymes in various tissues of male Fischer 344 rats are altered by food restriction. The Journal of nutrition 1995, 125(2):195–201.

69. Ivashchenko O, Van Veldhoven PP, Brees C, Ho YS, Terlecky SR, Fransen M: Intraperoxisomal redox balance in mammalian cells: oxidative stress and interorganellar cross-talk. Molecular biology of the cell 2011, 22(9):1440–1451.

70. Perichon R, Bourre JM: Peroxisomal beta-oxidation activity and catalase activity during development and aging in mouse liver. Biochimie 1995, 77(4):288–293.

71. Weir HJ, Yao P, Huynh FK, Escoubas CC, Goncalves RL, Burkewitz K, Laboy R, Hirschey MD, Mair WB: Dietary Restriction and AMPK Increase Lifespan via Mitochondrial Network and Peroxisome Remodeling. Cell Metab 2017, 26(6):884–896 e885.

72. Houtkooper RH, Argmann C, Houten SM, Canto C, Jeninga EH, Andreux PA, Thomas C, Doenlen R, Schoonjans K, Auwerx J: The metabolic footprint of aging in mice. Scientific reports 2011, 1:134.

73. Poirier Y, Antonenkov VD, Glumoff T, Hiltunen JK: Peroxisomal beta-oxidation--a metabolic pathway with multiple functions. Biochimica et biophysica acta 2006, 1763(12):1413–1426.

74. Singh J, Khan M, Singh I: Silencing of Abcd1 and Abcd2 genes sensitizes astrocytes for inflammation: implication for X-adrenoleukodystrophy. Journal of lipid research 2009, 50(1):135–147.

75. Pujol A, Hindelang C, Callizot N, Bartsch U, Schachner M, Mandel JL: Late onset neurological phenotype of the X-ALD gene inactivation in mice: a mouse model for adrenomyeloneuropathy. Human molecular genetics 2002, 11(5):499–505.

76. Seedorf U, Raabe M, Ellinghaus P, Kannenberg F, Fobker M, Engel T, Denis S, Wouters F, Wirtz KW, Wanders RJ et al: Defective peroxisomal catabolism of branched fatty acyl coenzyme A in mice lacking the sterol carrier protein-2/sterol carrier protein-x gene function. Genes & development 1998, 12(8):1189–1201.

77. Im SS, Yousef L, Blaschitz C, Liu JZ, Edwards RA, Young SG, Raffatellu M, Osborne TF: Linking lipid metabolism to the innate immune response in macrophages through sterol regulatory element binding protein-1a. Cell Metab 2011, 13(5):540–549.

78. Di Cara F, Sheshachalam A, Braverman NE, Rachubinski RA, Simmonds AJ: Peroxisome-Mediated Metabolism Is Required for Immune Response to Microbial Infection. Immunity 2017, 47(1):93–106 e107.

79. Li Z, Li J, Bu X, Liu X, Tankersley CG, Wang C, Huang K: Age-induced augmentation of p38 MAPK phosphorylation in mouse lung. Experimental gerontology 2011, 46(8):694–702.

80. Kim HJ, Jung KJ, Yu BP, Cho CG, Chung HY: Influence of aging and calorie restriction on MAPKs activity in rat kidney. Experimental gerontology 2002, 37(8-9):1041–1053.

81. Kim EK, Choi EJ: Pathological roles of MAPK signaling pathways in human diseases. Biochimica et biophysica acta 2010, 1802(4):396–405.

82. Vrailas-Mortimer A, del Rivero T, Mukherjee S, Nag S, Gaitanidis A, Kadas D, Consoulas C, Duttaroy A, Sanyal S: A muscle-specific p38 MAPK/Mef2/MnSOD pathway regulates stress, motor function, and life span in Drosophila. Developmental cell 2011, 21(4):783–795.

83. Wang MC, Bohmann D, Jasper H: JNK signaling confers tolerance to oxidative stress and extends lifespan in Drosophila. Developmental cell 2003, 5(5):811–816.

84. Oh SW, Mukhopadhyay A, Svrzikapa N, Jiang F, Davis RJ, Tissenbaum HA: JNK regulates lifespan in Caenorhabditis elegans by modulating nuclear translocation of forkhead transcription factor/DAF-16. Proc Natl Acad Sci U S A 2005, 102(12):4494–4499.

85. Biteau B, Karpac J, Hwangbo D, Jasper H: Regulation of Drosophila lifespan by JNK signaling. Experimental gerontology 2011, 46(5):349–354.

86. Johnson GL, Nakamura K: The c-jun kinase/stress-activated pathway: regulation, function and role in human disease. Biochimica et biophysica acta 2007, 1773(8):1341–1348.

87. Sun J, Kale SP, Childress AM, Pinswasdi C, Jazwinski SM: Divergent roles of RAS1 and RAS2 in yeast longevity. J Biol Chem 1994, 269(28):18638–18645.

88. Slack C, Alic N, Foley A, Cabecinha M, Hoddinott MP, Partridge L: The Ras-Erk-ETS-Signaling Pathway Is a Drug Target for Longevity. Cell 2015, 162(1):72–83.

89. Nanji M, Hopper NA, Gems D: LET-60 RAS modulates effects of insulin/IGF-1 signaling on development and aging in Caenorhabditis elegans. Aging Cell 2005, 4 (5):235–245.

90. Thyagarajan B, Blaszczak AG, Chandler KJ, Watts JL, Johnson WE, Graves BJ: ETS-4 is a transcriptional regulator of life span in Caenorhabditis elegans. PLoS Genet 2010, 6(9):e1001125.

91. Owusu-Ansah E, Yavari A, Banerjee U: A protocol for in vivo detection of reactive oxygen species. Protocol Exchange 2008.

92. Afgan E, Baker D, van den Beek M, Blankenberg D, Bouvier D, Cech M, Chilton J, Clements D, Coraor N, Eberhard C et al: The Galaxy platform for accessible, reproducible and collaborative biomedical analyses: 2016 update. Nucleic Acids Res 2016, 44(W1):W3–W10.

93. Kim D, Pertea G, Trapnell C, Pimentel H, Kelley R, Salzberg SL: TopHat2: accurate alignment of transcriptomes in the presence of insertions, deletions and gene fusions. Genome biology 2013, 14(4):R36.

94. Trapnell C, Roberts A, Goff L, Pertea G, Kim D, Kelley DR, Pimentel H, Salzberg SL, Rinn JL, Pachter L: Differential gene and transcript expression analysis of RNA-seq experiments with TopHat and Cufflinks. Nature protocols 2012, 7(3):562–578.

95. Love MI, Huber W, Anders S: Moderated estimation of fold change and dispersion for RNA-seq data with DESeq2. Genome biology 2014, 15(12):550.

96. Subramanian A, Tamayo P, Mootha VK, Mukherjee S, Ebert BL, Gillette MA, Paulovich A, Pomeroy SL, Golub TR, Lander ES et al: Gene set enrichment analysis: a knowledge-based approach for interpreting genome-wide expression profiles. Proc Natl Acad Sci U S A 2005, 102(43):15545–15550.

97. Pfaffl MW, Tichopad A, Prgomet C, Neuvians TP: Determination of stable housekeeping genes, differentially regulated target genes and sample integrity: BestKeeper––Excel-based tool using pair-wise correlations. Biotechnology letters 2004, 26(6):509–515.

98. Schmittgen TD, Livak KJ: Analyzing real-time PCR data by the comparative C(T) method. Nature protocols 2008, 3(6):1101–1108.

